# Natural variability in bee brain size and symmetry revealed by micro-CT imaging and deep learning

**DOI:** 10.1101/2022.10.12.511944

**Authors:** Philipp D. Lösel, Coline Monchanin, Renaud Lebrun, Alejandra Jayme, Jacob Relle, Jean-Marc Devaud, Vincent Heuveline, Mathieu Lihoreau

## Abstract

Analysing large numbers of brain samples can reveal minor, but statistically and biologically relevant variations in brain morphology that provide critical insights into animal behaviour, ecology and evolution. So far, however, such analyses have required extensive manual effort, which considerably limits the scope for comparative research. Here we used micro-CT imaging and deep learning to perform automated analyses of 3D image data from 187 honey bee and bumblebee brains. We revealed strong inter-individual variations in total brain size that are consistent across colonies and species, and may underpin behavioural variability central to complex social organisations. In addition, the bumblebee dataset showed a significant level of lateralization in optic and antennal lobes, providing a potential explanation for reported variations in visual and olfactory learning. Our fast, robust and user-friendly approach holds considerable promises for carrying out large-scale quantitative neuroanatomical comparisons across a wider range of animals. Ultimately, this will help address fundamental unresolved questions related to the evolution of animal brains and cognition.

**Author Summary:** Bees, despite their small brains, possess a rich behavioural repertoire and show significant variations among individuals. In social bees this variability is key to the division of labour that maintains their complex social organizations, and has been linked to the maturation of specific brain areas as a result of development and foraging experience. This makes bees an ideal model for understanding insect cognitive functions and the neural mechanisms that underlie them. However, due to the scarcity of comparative data, the relationship between brain neuro-architecture and behavioural variance remains unclear. To address this problem, we developed an AI-based approach for automated analysis of brain images and analysed an unprecedentedly large dataset of honey bee and bumblebee brains. Through this process, we were able to identify previously undescribed anatomical features that correlate with known behaviours, supporting recent evidence of lateralized behaviour in foraging and pollination. Our method is open-source, easily accessible online, user-friendly, fast, accurate, and robust to different species, enabling large-scale comparative analyses across the animal kingdom. This includes investigating the impact of external stressors such as environmental pollution and climate change on cognitive development, helping us understand the mechanisms underlying the cognitive abilities of animals and the implications for their survival and adaptation.

## Introduction

Artificial intelligence is helping scientists to more efficiently and effectively analyse data in a wide range of scientific fields, enabling them to make new discoveries and address important open questions [1]. In particular, neuroscience is a field that can greatly benefit from automated analysis tools for large-scale comparative investigations. Animals, from insects to humans, show a rich diversity of behavioural profiles, or personalities, that may be underpinned by anatomical and cognitive variability [2]. Identifying natural variations in brain morphology and performance at the intra- and inter-specific levels can therefore help to understand the evolution of species and their potential for resilience to environmental stressors in the context of biodiversity loss [3]. However, such an approach currently requires extensive manual effort in order to obtain and analyse large and high-quality brain datasets, limiting investigations to a few individuals and model species for which brain atlases are available (e.g. rodents [4], primates [5], *Drosophila* [6]), thereby restraining the scope for comparative research [7].

Three-dimensional (3D) imaging techniques, such as micro-computed tomography (micro-CT), offer the potential to facilitate large-scale investigations. These methods enable non-destructive, fine-scale imaging of internal structures of biological objects, including brain tissues, both *in vivo* and *ex vivo.* Recent improvements in resolution and acquisition times [8,9] have broadened the application of micro-CT to a wide range of animals, including non-model species of small sizes like insects (e.g. ants [10], wasps [11], beetles [12], bees [13,14]), and posed new demands for accelerated image analysis methods. However, image segmentation and post-processing steps still require manual assistance [15–18], which is time-consuming (several hours or days to reconstruct a structure) and not feasible for analysing large datasets.

Here we demonstrate how deep learning can significantly speed up 3D image analysis for large-scale comparative brain studies in non-model species by using the semi-automated [19,20] and automated segmentation methods of the recently developed online platform Biomedisa [21] (https://biomedisa.org) to analyse micro-CT image data from bee brains (Fig 1).

**Fig 1.**
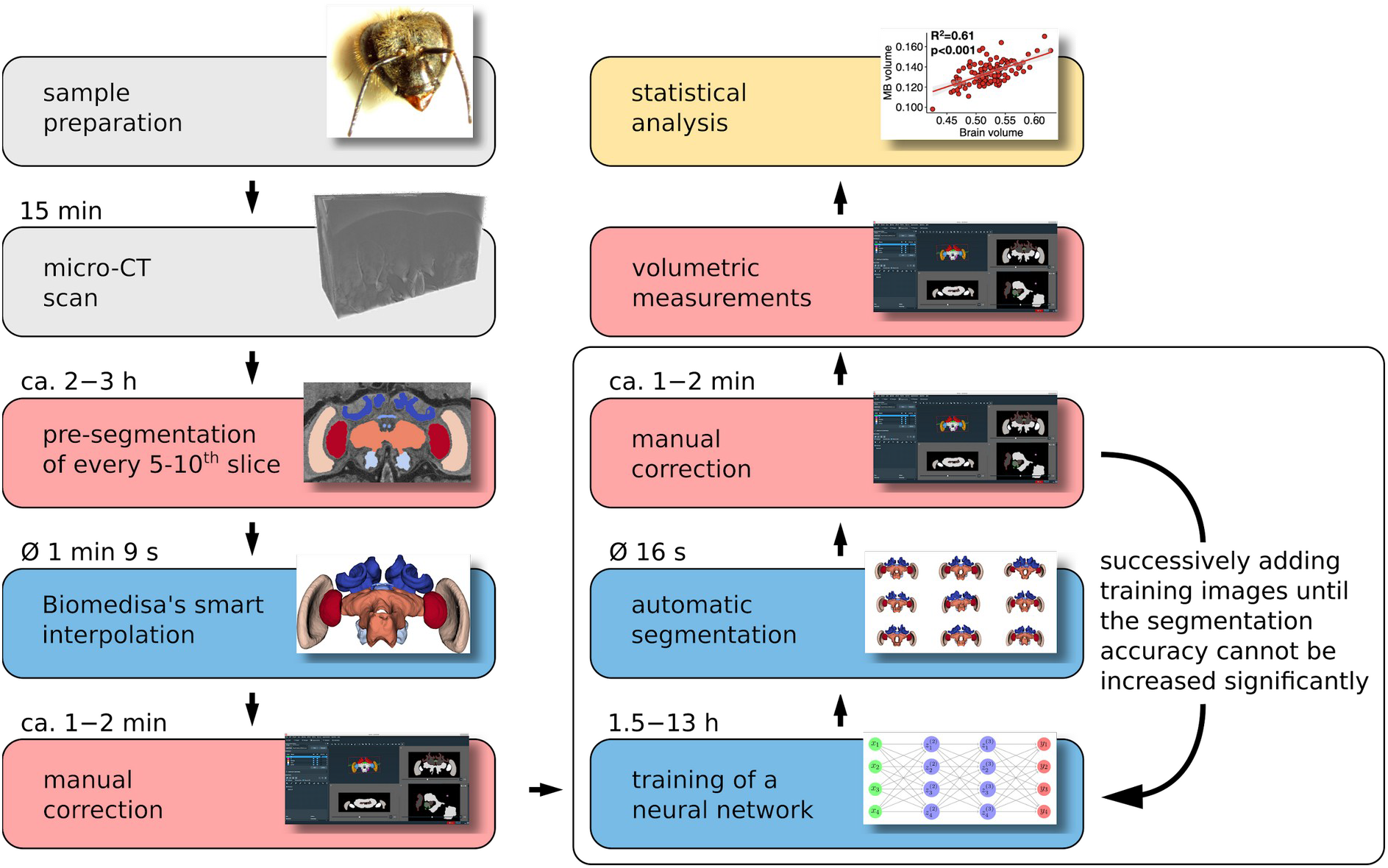
Flowchart of the steps to perform large-scale quantitative comparative analyses of bee brain size and organisation using micro-CT imaging and the Biomedisa segmentation platform. After sample preparation and volume reconstruction (grey boxes), the micro-CT scans are segmented with AVIZO 2019.1 (red boxes) in combination with Biomedisa (blue boxes). The volumes are then measured with AVIZO 2019.1 and statistically analysed with R Studio (yellow box).

Despite their small size (<1 mm^3^), bee brains support a rich behavioural repertoire and exhibit extensive inter-individual variability [22]. In social bees, such as honey bees and bumblebees, this variability is central to the division of labour that supports complex social organisations, and some of it has been linked to the maturation of specific brain areas due to development and foraging experience [23,24]. This makes bees ideal models for studying insect cognitive functions and their underlying neural substrates [25,26]. However, there is currently no clear connection between possible differences in brain neuro-architecture and behavioural variance due to the lack of data from comparative analyses (few studies [27] and sample sizes ranging from 7 [28] to 67 [29] individuals).

By collecting volumetric data on brains of 110 honey bees and 77 bumblebees and using semi-automated segmentation to annotate the training data for a deep neural network, we were able to reduce data processing time by up to 98% compared to a conventional manual segmentation approach. Our results revealed size variations and strong anatomical asymmetries in the brain samples, supporting behavioural observations from the literature. In addition to an extensive evaluation and a detailed protocol of Biomedisa’s deep learning method, and the new insights into the evolution of bee brains, this is the first study to our knowledge that will publish an unprecedentedly large dataset of the underlying image and label data of bee brains. Our method is open access, readily available online, fast, accurate, robust to different species, and can be applied to a wide range of 3D imaging modalities and scientific questions beyond comparative neurosciences.

## Results and Discussion

### Automatic segmentation of bee brains considerably speeds up analysis

We performed micro-CT scanning of 120 honey bee foragers (*Apis mellifera*, Buckfast) from two apiaries (Population A: 100 bees from 6 hives, population B: 20 bees from 3 hives) in Toulouse (France). Brain samples were prepared following Smith et al. [14] and CT-scanned at a resolution of 5.4 μm isotropic voxel size (see “Methods”). Among the 120 micro-CT scanned brains, 10 were damaged during manipulation and discarded, leaving 110 brains for our analysis. Image dimensions and image spacing varied across subjects and averaged 844 × 726 × 485 isotropic voxels and 0.0054 × 0.0054 × 0.0054 mm^3^, respectively.

We analysed six major brain neuropils based on the 3D bee brain atlas [30]: the antennal lobes (AL) that receive olfactory information from the antennae; the mushroom bodies (MB, each comprising the medial and lateral calyces, peduncle and lobe) that integrate olfactory and visual information; the central complex (CX, comprising the central body, the paired noduli and the protocerebral bridge) that receives compass and optic flow information; the medullae (ME) and lobulae (LO) that receive visual information from the compound eyes, combined together as ‘optic lobes’ (OL) in our analysis (retinae and laminae were not measured); and the other remaining neuropils (OTH) (including protocerebral lobes and subesophageal zone) (Fig 2). ALs and OLs are involved in olfactory and visual processing, respectively, while the CX and MBs play important roles in locomotor behaviour, learning and memory, respectively [31].

**Fig 2.**
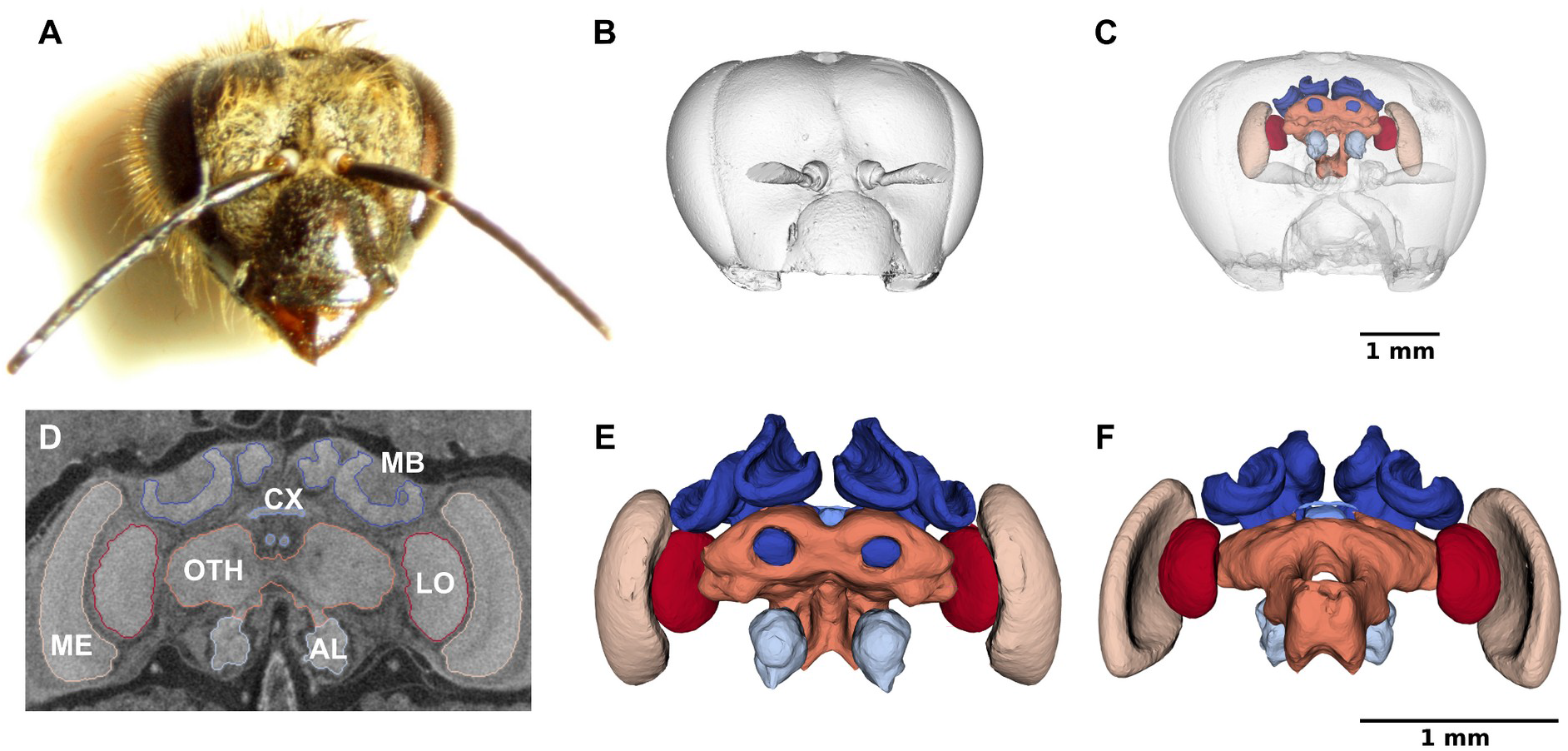
Surface renderings of an example of CT-scanned honey bee head and reconstructed brain neuropils. (**A**) Frontal view of the head of a forager bee (ID 79, hive H4). (**B**) Surface rendering of the head with the mandibles removed. (**C**) Overlay of the head and reconstructed neuropils. (**D**) Frontal cross-section of the tomogram with the segmentation boundaries of the mushroom bodies (MB), central complex (CX), antennal lobes (AL), medullae (ME), lobulae (LO) and other neuropils (OTH). (**E**) Frontal view of the reconstructed MB (*dark blue*), CX (*sky blue*), AL (*light sky blue*), ME (*beige*), LO (*red*) and OTH (*orange*). (**F**) Dorsal view of the reconstructed neuropils. (B), (C), (E) and (F) were created with ParaView Glance integrated in Biomedisa.

The volumetric analysis of the different neuropils required their isolation from the CT scans by segmentation (Figs 1 and 2). Each dataset was manually cropped to the area of the neuropils (Fig 2D) using AVIZO 2019.1, resulting in an average size of 451 × 273 × 167 voxels. For automatic segmentation, we used the deep neural network interface from the online platform Biomedisa. To train a deep neural network, Biomedisa needs a set of fully segmented volumetric images. To create the first three datasets, labels were assigned manually to the six neuropils in every 10th slice and every 5th slice in the interval containing CX within the 3D volume using AVIZO 2019.1 (Thermo Fisher Scientific, Waltham, USA). For better performance compared to purely morphological interpolation (see “Methods” and Fig 3), Biomedisa’s semi-automatic smart interpolation was used to segment the remaining volume between the pre-segmented slices. Before interpolation, the image data was slightly smoothed using Biomedisa’s ‘denoise’ function (see “Methods”). Subsequently, outliers (i.e. unconnected voxels or islands) were removed and segmentation errors were corrected manually by an expert using AVIZO 2019.1.

**Fig 3.**
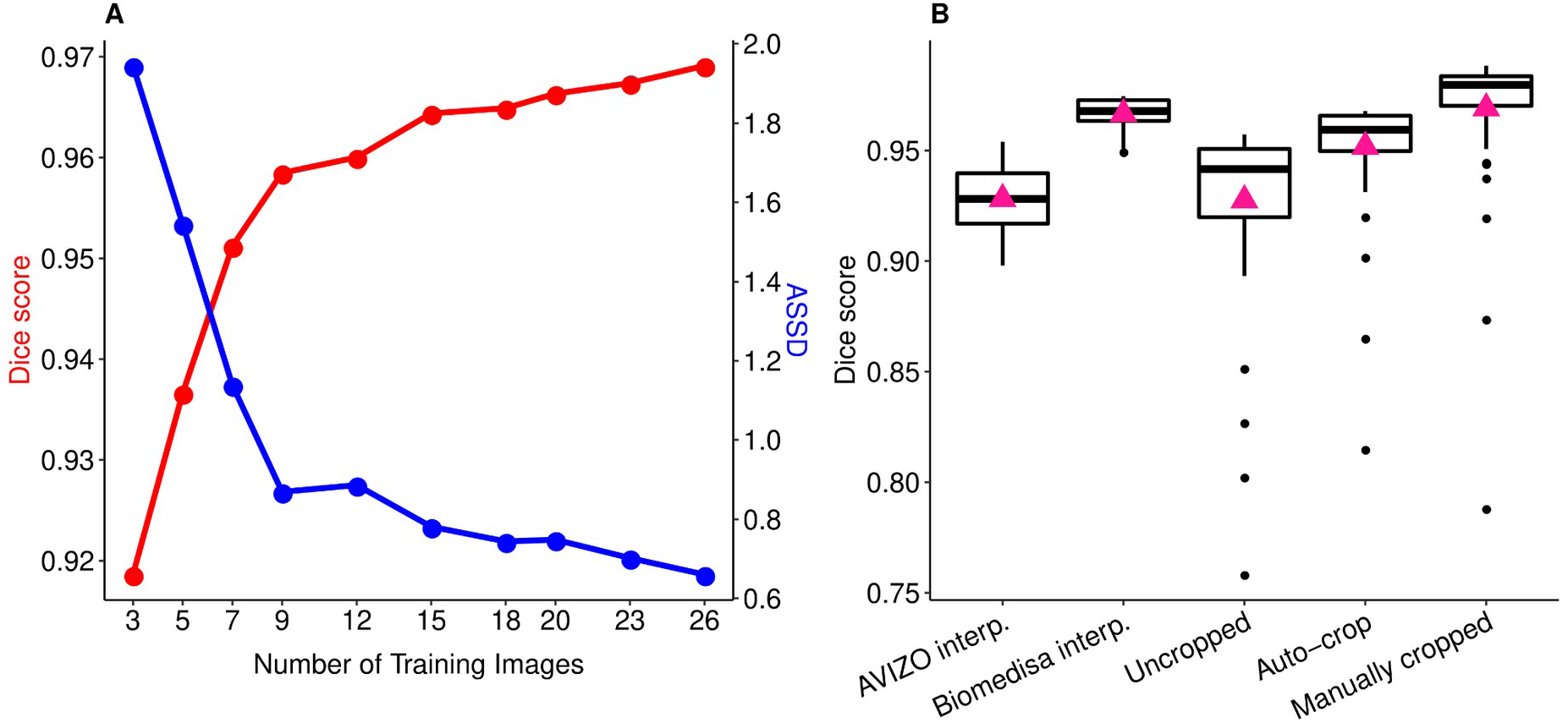
Segmentation accuracy of Biomedisa’s semi-automatic and automatic segmentation of honey bee CT scans. (**A**) Average Dice scores and ASSD of the automatic segmentation results for an increasing number of training images. (**B**) Semi-automated segmentation accuracy (Dice score) of the AVIZO interpolation and the Biomedisa interpolation as well as segmentation accuracy of the automatic segmentation for uncropped image data, using Biomedisa’s auto-copping and manually cropped image data. Boxplots show median volumes (intermediate line) and quartiles (upper and lower lines). Pink triangles display mean volumes. For performance tests of the automatic segmentation, the 84 honey bee test images were split into 30 validation images and 54 test images (see “Methods”).

Additional training data was then iteratively created using neural networks. Starting from three semi-automatically segmented images, we trained neural networks on 3, 7, 12, 18, and 26 images by adding manually corrected segmentation results of the last trained network to the training data after each step (Fig 1). In all cases, the network’s default configuration was used (see “Methods”). The corresponding training times on 4 NVIDIA Tesla V100s were 1.5, 3.5, 6, 9, and 13 hours. At this point, segmentation accuracy could no longer be significantly improved by additional training data (Fig 3 and S1 Table). We finally used the network trained on the set of 26 images (“training images”) to automatically segment the remaining 84 micro-CT scans of honey bee brains (“test images”). The automatic segmentation of each 3D image took an average of 16 seconds using an NVIDIA RTX A5000 or 82 seconds using an Intel^®^ Core^™^ i9-11900K Processor. All segmentation results were checked and manually corrected by an expert (using ‘remove islands’ and ‘merge neighbours’ functions in AVIZO 2019.1).

To test the suitability of our approach for brain datasets from other species (that, unlike honey bees [26], are not well-established model species in neurosciences), we also performed micro-CT scans of 77 bumblebees (*Bombus terrestris*) from 4 commercially-available colonies (colony A: 18 bumblebees, colony B: 20 bumblebees, colony C: 21 bumblebees, colony D: 18 bumblebees). Based on the evaluation for an increasing number of training images (see “Methods”, Fig 3 and S1 Table), we decided to semi-automatically segment 13 training images (which we considered to be a good balance between manual effort and accuracy) using Biomedisa to train a neural network on the bumblebee dataset. The trained network was then used to automatically segment the remaining 64 CT scans.

Overall, AVIZO 2019.1 was used for pre-segmentation, correction of segmentation results and measurement of absolute neuropil volumes calculated with the voxel count function. Biomedisa was used for smart interpolation to create the initial training data, training of deep neural networks and subsequent automatic segmentation. Since Biomedisa supports the AMIRA Mesh File format, deployed by AVIZO and AMIRA, data can be easily transferred between the Biomedisa online platform and AVIZO 2019.1. In addition to AMIRA meshes (AM), Biomedisa supports many other common data formats (such as DICOM, NifTI, TIFF, NRRD, MHA, MHD) and can also be used in combination with other segmentation tools.

For both datasets, this automatic segmentation considerably reduced the time and effort required for image analysis compared to purely manual segmentation. For each 3D image, it took about 5 to 10 minutes for the whole procedure: importing the data into AVIZO 2019.1, cropping to the area of the neuropils, exporting the data, uploading it to Biomedisa, performing automatic segmentation, downloading and importing the segmentation result into AVIZO 2019.1, correcting the segmentation result manually, and measuring volumes. In contrast, a purely manual or semiautomatic approach would have taken several hours for each CT scan (approx. 5 to 6 h using conventional manual segmentation and 2 to 3 h using Biomedisa’s semi-automatic segmentation). Depending on the quality of the automated segmentation result, either no correction was required (S1A Fig) or manual correction typically took 1 to 2 minutes (S1B Fig), rarely longer if the result was significantly flawed (S1C Fig). Typical artefacts resulting from the automatic segmentation are outliers (S1B Fig) which can be easily removed either with Biomedisa’s cleaning function or with AVIZO 2019.1. In the end, with our approach, the manual effort for segmenting the full honey bee dataset (15 to 27 h) was about 95 to 98% less than purely manual segmentation (550 to 660 h) and about 87 to 93% less for the bumblebee dataset (31 to 50 h compared to 385 to 462 h) (see “Methods”).

### Automatic segmentation of bee brains is highly accurate

We evaluated the accuracy of the automatic segmentation using two complementary and commonly applied metrics for measuring performance in biomedical image segmentation challenges [32] (see “Methods”): the Dice similarity coefficient (Dice) and the average symmetric surface distance (ASSD). The Dice score quantifies the matching level between two segmentations and is between 0 (no overlap at all) and 1 (perfect match). The ASSD is the average 3D Euclidean distance from a point on the surface of one segmentation to the closest point on the surface of the second segmentation and vice versa. The smaller the ASSD, the closer the two segmentations.

For both bee species, we measured the accuracy of the segmentation results of the test images performed by the deep neural network trained with the respective training data. We compared the results obtained without further manual post-processing with ground truth data generated by an expert (i.e. segmentation results after revision and manual correction). For the automatic segmentation of the 84 honey bee brains, a Dice score of 0.988 was achieved (S1 Table). In 10.7% of the test images, little or no manual correction was necessary (error less than 0.01% of Dice score, S1A Fig). In 84.5%, a slight manual correction was required taking 1 to 2 minutes (error ranging between 0.01% and 4%, S1B Fig). Only 4.8% of the segmentation results were significantly flawed (error greater than 4%, S1C Fig), usually due to a significant deviation from the training data (e.g. images from incompletely stained or damaged brains during tissue processing) and required extensive manual correction or semi-automatic reconstruction using Biomedisa’s smart interpolation. For the automatic segmentation of the 64 bumblebee brains, a Dice score of 0.983 was achieved (S1 Table). Here, no error was less than 0.01%. However, for 9.4% of the segmentation results, the error was less than 0.1%. For 78.1% it was between 0.1% and 4% and 12.5% had an error greater than 4%.

Biomedisa thus enabled high-precision and fast segmentation of the 148 test images (84 honey bees, 64 bumblebees), which required manual correction of the segmentation results equalling a Dice score of 0.012 (i.e. an error of 1.2%) for the honey bee brains and 0.017 for the bumblebee brains (S1 Table). Note that major corrections were mostly only required for the smallest brain area considered, the CX (an average error of 3.3% for honey bees and 16.3% for bumblebees), because its fine structure combined with low contrast often makes it difficult to detect this neuropil, particularly in the bumblebee CT scans.

### Brain volumes varied significantly among honey bees

To validate our segmentation approach, we compared our results with previously published data, focusing on the volumes of the six neuropils considered (Fig 4) in honey bees. Division of labour in honey bee colonies is primarily based on age differences among individuals [33]. Since workers vary little in body size (34], it was expected that their brain volume would also show only small variation. Our results are fairly consistent with previous studies based on smaller datasets [24,28–30,35–41] and using different quantitative approaches (CT scan data [40,41], or stereo [24,28,29,37], confocal [30,35,36], and nuclear magnetic resonance [35] microscopy). When data do not match, we expect our measures to be more accurate than those in previous studies since our measurements were taken from thinner slices (5.4 μm vs. 8 [30] to 60 [38] μm) and on all successive slices (rather than leaving intermediate slices aside and using linear interpolation, e.g. Cavalieri principle [42]). Such technical discrepancies likely explain under- or over-estimated volumes in other studies compared to ours. Also note that some measurements from other studies have uncertainties, e.g. a precise description of which structures were included in the measurement of CX [30] or whether or not cell bodies were included in the total brain volume [28]. Additionally, differences in the estimated structural volumes may arise from biological differences among samples taken from bees of different subspecies (e.g. European and Africanized honey bees [29]), ages and foraging experiences of individual bees [24,39].

**Fig 4.**
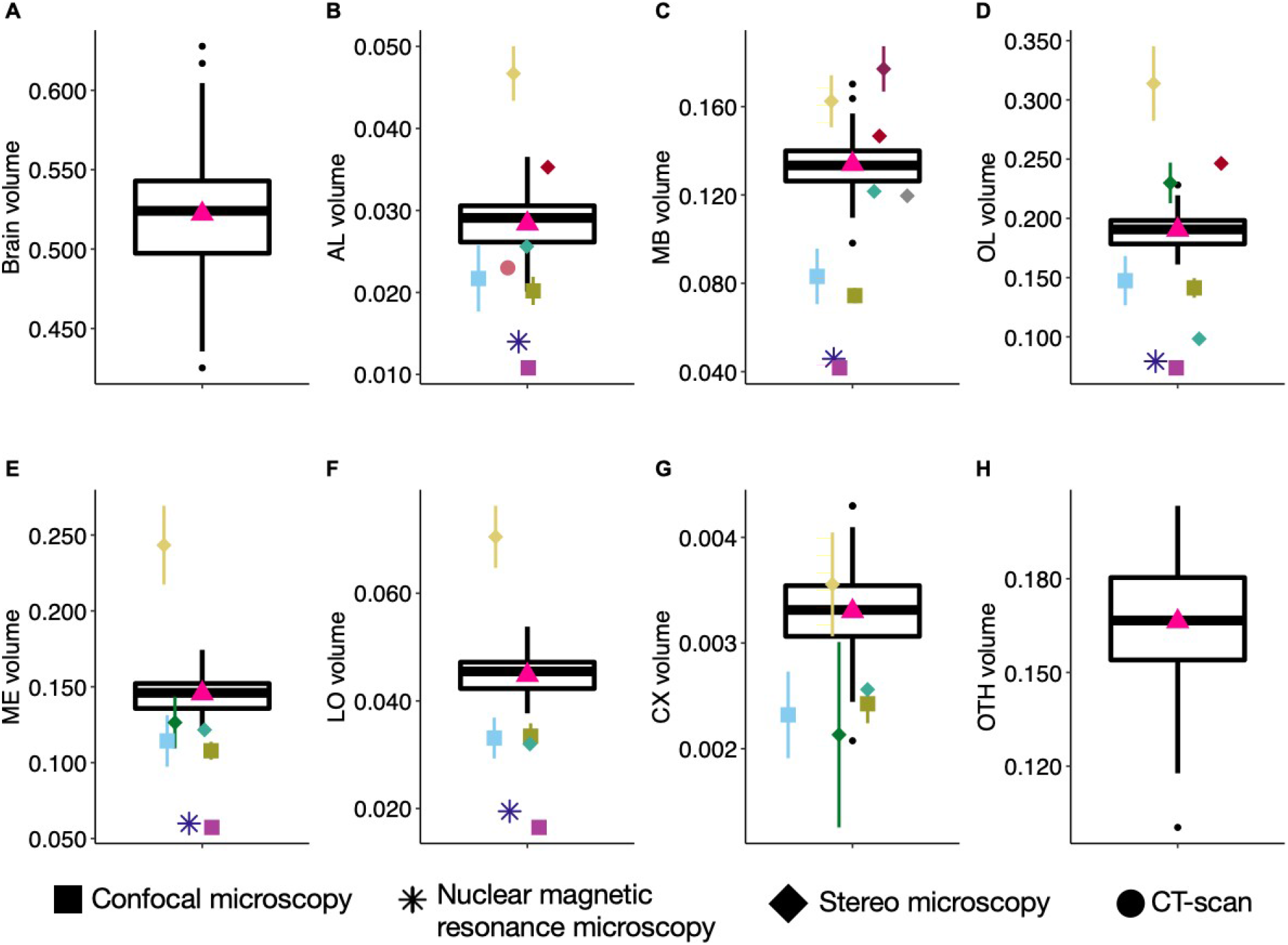
Variation in brain and neuropils volumes (mm^3^) for honey bees (N=110). (**A**) Total brain. (**B**) Antennal lobes (AL). (**C**) Mushroom bodies (MB). (**D**) Optic lobes (OL). (**E**) Medullae (ME). (**F**) Lobulae (LO). (**G**) Central complex (CX). (**H**) Other neuropils (OTH). Boxplots show median volumes (intermediate line) and quartiles (upper and lower lines). Pink triangles display mean volumes. Coloured symbols show mean (± s.d. when available) of neuropil volumes described for forager honey bees in other studies: using confocal microscopy (*square*): Brandt et al. [30] (N=20 bees - *light blue*); Steijven et al. [36] (N=10 - *kaki*); Haddad et al. [35] (*purple);* using nuclear magnetic resonance microscopy (*star*): Haddad et al. [35] (N=8 - *deep blue*); using stereo microscopy (*diamond):* Gowda & Gronenberg [28] (N=7 - *yellow*); Gronenberg & Couvillon [29] (N=121 European and Africanized honey bees - *dark green*); Mares et al. [37] (N=25 - *turquoise*); Maleszka et al. [38] (N=30 - *burgundy*);Withers et al. [24] (*red*); Durst et al. [39] (N=12 - *grey*); using CT scan (*point*): Greco and Stait-Gardner [40] (N=10 - *coral*). For the total brain volume, comparisons with other studies are not shown because of the cumulative uncertainties of the measurements.

Among our 110 honey bee brains, total brain volume (i.e. sum of all measured neuropils) varied by 32% and neuropil volumes by 29 to 52% (Fig 4 and Table 1). We found strong positive correlations between the absolute volumes of all neuropils - but the CX - and total brain volume (Figs 5A–G). Most neuropils scaled isometrically with total brain volume (Fig 5H), with only a lower correlation coefficient for CX (r=0.36, p<0.001; Fig 5F). This is coherent with the results of the largest comparative analysis of honey bee brains so far (121 brain samples obtained by electron microscopy technique in two honey bee strains) [29]. When considering relative volumes (ratio of neuropil volume to total brain volume), we found only a positive correlation between OTH and total brain volumes, while relative volumes of MB, OL, ME and CX correlated negatively with total brain volume (S2G Fig).

**Table 1.**
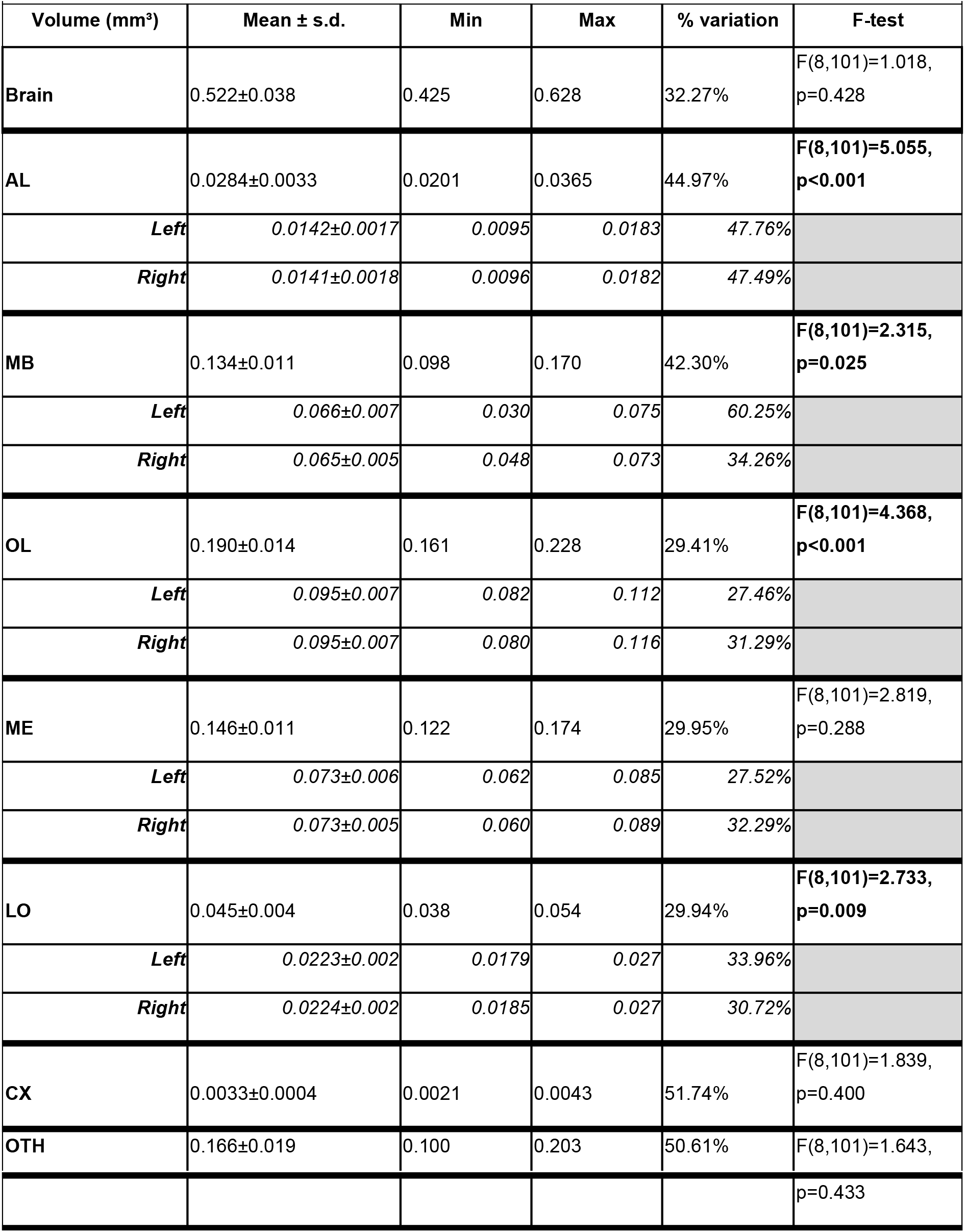
Total brain and neuropil volumes of honey bees (N=110). Mean (± standard deviation), minimal and maximal volumes (mm^3^) and percentage of volume variation ((Max-Min)*100/Max). For paired neuropils, detailed data for both sides (left/right) are given. Note that the left/right comparison for MB is based on data from 59 honey bees only, as some bees have both sides merged in the automatic segmentation result. F-test, following LMMs, tests the significance of the fixed variable ‘hive’, and results are displayed in bold when significant.

**Fig 5.**
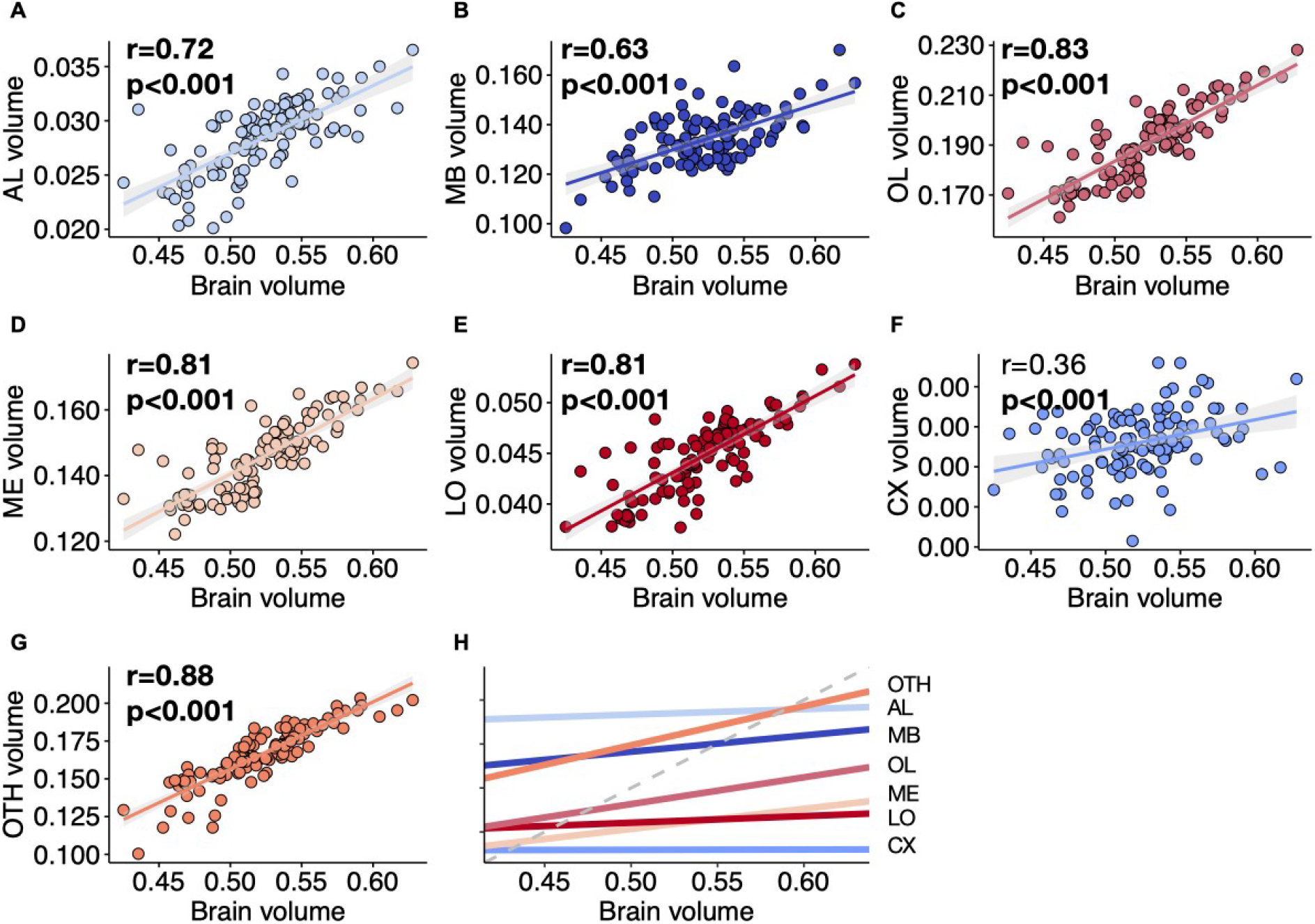
Correlation between neuropil volumes and total brain volume (mm^3^) (N=110 honey bees). (**A**) Antennal lobes (AL). (**B**) Mushroom bodies (MB). (**C**) Optic lobes (OL). (**D**) Medullae (ME). (**E**) Lobulae (LO). (**F**) Central complex (CX). (**G**) Other neuropils (OTH). Regression lines displayed with 95% confidence intervals. Pearson correlation coefficient (r) and p-value are given. Strong correlations (r>0.40) and significant correlations (p<0.05) are displayed in bold. (**H**) Linear correlations for the different neuropils (y-axis not given: differs for each neuropil). The grey dashed line indicates true isometric correlation (slope=1).

When assessing correlations between absolute volumes across neuropils, we found strong correlations between OL, LO and ME volumes (S2 Table), as previously reported [29]. In our dataset, however, AL volume is positively correlated with all other neuropils. We also found positive correlations for MB, CX and OTH with all other neuropils, except between MB and CX, which were not correlated. We next compared the relative neuropil volumes in order to search for relevant allometric relationships (S2 Table). Again, we found strong correlations between the relative volumes of OL, LO and ME, consistent with previous studies [29]. There was a negative correlation between the relative volumes of AL and MB, while a previous study [29] reported a positive correlation between the relative volumes of AL and MB lobes (but not the calyces), but in our study we did not distinguish the structures within MBs. In addition, the relative volume of AL was positively correlated with those of OL, ME, and LO, while previous work on honey bees [29] and other bee species [43] suggested a negative correlation between AL and OL as a possible trade-off between visual and olfactory processing. Also, the relative OTH volume was negatively correlated with all other neuropils except AL. Overall, honey bees with larger brains tend to have larger neuropils, with the exception of CX, which is not as strongly related to brain size as the other neuropils. This weaker correlation could also be due to the small size of the neuropil and potential measure uncertainties caused by the difficulty of its segmentation.

### Variance in honey bee brain volumes was similar within and between colonies

Next, we explored the distribution of honey bee brain volumes within and between colonies. Social insect colonies can be considered superorganisms, characterised by a division of labour among workers that is partly determined by the genetics, age and morphology of individuals [44]. In honey bees, MB volume increases with age [24] and foraging experience [39,45], and AL volume correlates with behavioural tasks, i.e. nurse, comb-builder, nectar or pollen forager [46]. These variations are thought to support division of labour, as larger volumes (relative and/or absolute) of specific neuropils may allow some individuals to be more efficient at certain tasks, or result from behavioural specialisations. For example, larger ALs and/or MBs may provide foragers with better abilities at learning spatial information and floral cues than in-hive workers [39]. While some studies investigated variations in brain size between species [28] or within species due to stress exposure (e.g. pesticides [13], nutrition [36]), to our knowledge, the magnitude of intra-and inter-colony variability under standard conditions has not yet been investigated. Stable within-hive variation in brain volumes across colonies with different genetic backgrounds and in different geographic areas could suggest selection for variability.

Honey bees in the nine colonies exhibited overall similar average brain volumes (F-test: p=0.428; Fig 6A and Table 1), hence, inter-colony differences do not explain the substantial (32%) inter-individual variability. Within colonies, the values were also relatively stable (S3 Table). However, several neuropils exhibited significant variations in their absolute volumes across colonies (Figs 6B-G and Table 1): AL (F-test: p<0.001; Fig 6B), MB (F-test: p=0.025; Fig 6C), OL (F-test: p<0.001; Fig 6D) and LO (F-test: p=0.009; Fig 6F). Interestingly, intra-colony variability of neuropils was generally lower and CX and OTH were the more variable (resp. 11 to 49% and 11 to 51%, depending on the colony), despite non-significant changes between colonies (S3 Table and Figs 6G–H). Notably, we only found low intra-colonial variations in MB volumes (7 to 30%), which are known to increase in size by 15% with age and foraging experience over the lifespan [24). This low level of variability is probably due to the fact that we only studied honey bee foragers (see “Methods”) and therefore could not reflect the reported brain plasticity between emerging workers, nurses and foragers.

**Fig 6.**
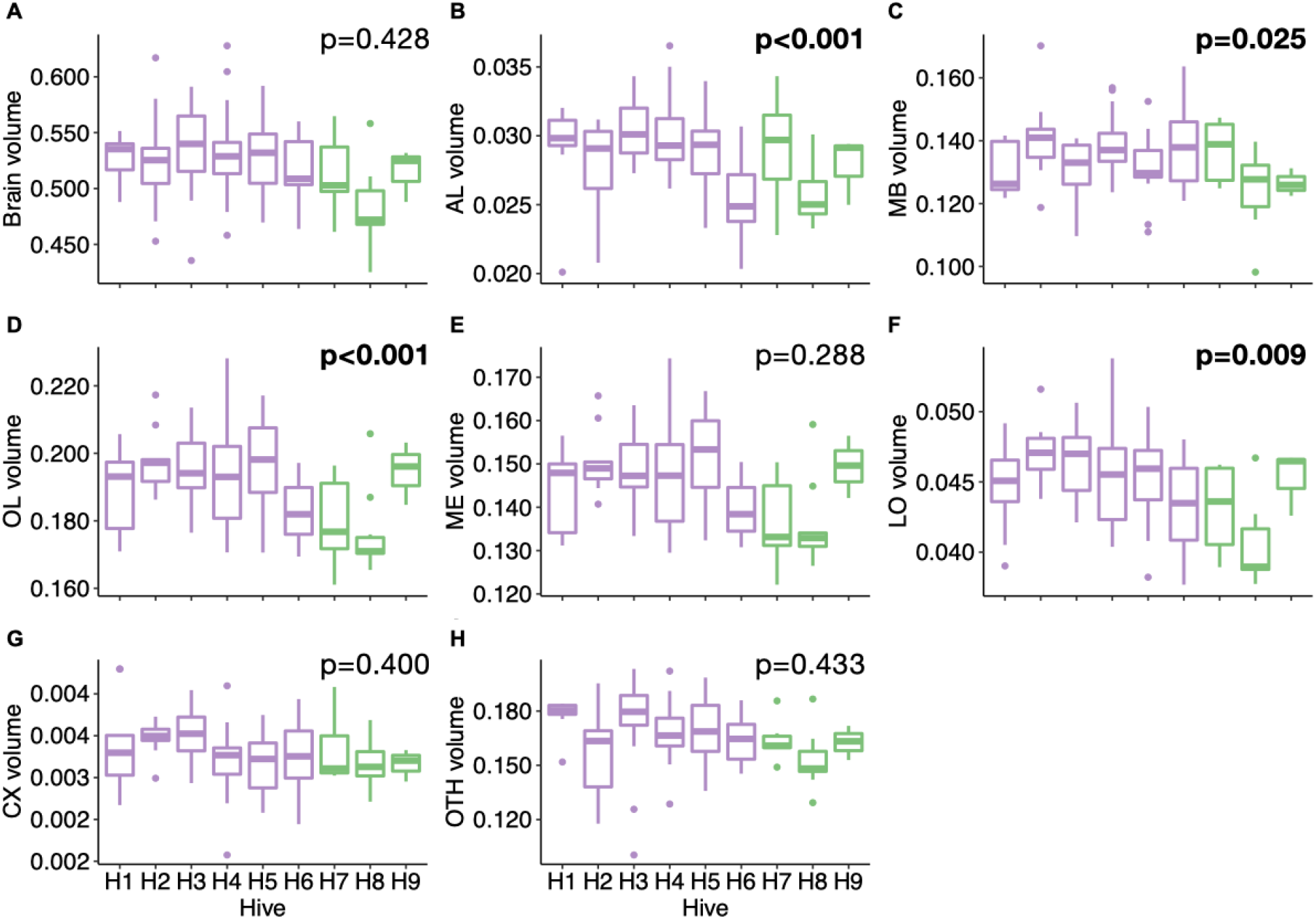
Variation in brain volume and absolute volume of neuropils (mm^3^) between honey bee colonies (population A - *purple*, N=6 hives; population B - *green*, N=3 hives). (**A**) Total brain. (**B**) Antennal lobes (AL). (**C**) Mushroom bodies (MB). (**D**) Optic lobes (OL). (**E**) Medullae (ME). (**F**) Lobulae (LO). (**G**) Central complex (CX). (**H**) Other neuropils (OTH). Statistical comparisons (p-values) for the neuropil volumes between hives were obtained from the F-test following LMMs and are displayed in bold when significant (statistical details in Table 1).

Thus, overall, we found moderate variability in brain size and neuropil volumes that were rather stable within colonies, based on a relatively homogeneous sample regarding strain (Buckfast) and behavioural specialisation (foragers). This suggests adaptive levels of brain size variation, potentially underpinning behavioural diversity in division of labour, particularly when specific brain areas are considered.

### Bumblebees showed higher variability in brain and neuropil volumes than honey bees

To demonstrate the generalisability of our approach to other sample series, and illustrate how this can be used to compare species, we applied the same analyses to the bumblebee brains. Bumblebees are increasingly used for comparative behavioural and cognitive research [47] and a brain atlas was recently published [48]. In contrast to honey bees, division of labour in bumblebees is primarily based on body size variation [49] with little effect of age (foragers tend to be larger than non-foragers [50]). Therefore, it is not surprising that overall bumblebees exhibited higher variation levels in total brain volume (52%; Fig 7A and S4 Table) and neuropil volumes (44 to 85%; Figs 7B–H), as compared to honey bees. Note, however, that the 77 bumblebees randomly sampled in the four source colonies were likely to be more heterogeneous than the 110 honey bees, which were only foragers. Yet, all neuropils scaled isometrically with total brain volume (Fig 7I), with again only a lower correlation for CX (r=0.59, p<0.001). Similar to honey bees, we found that only the relative volume of OTH was positively correlated with total brain volume and the relative volumes of LO, ME (and consequently OL) were negatively correlated with total brain volume (Fig 7J).

**Fig 7.**
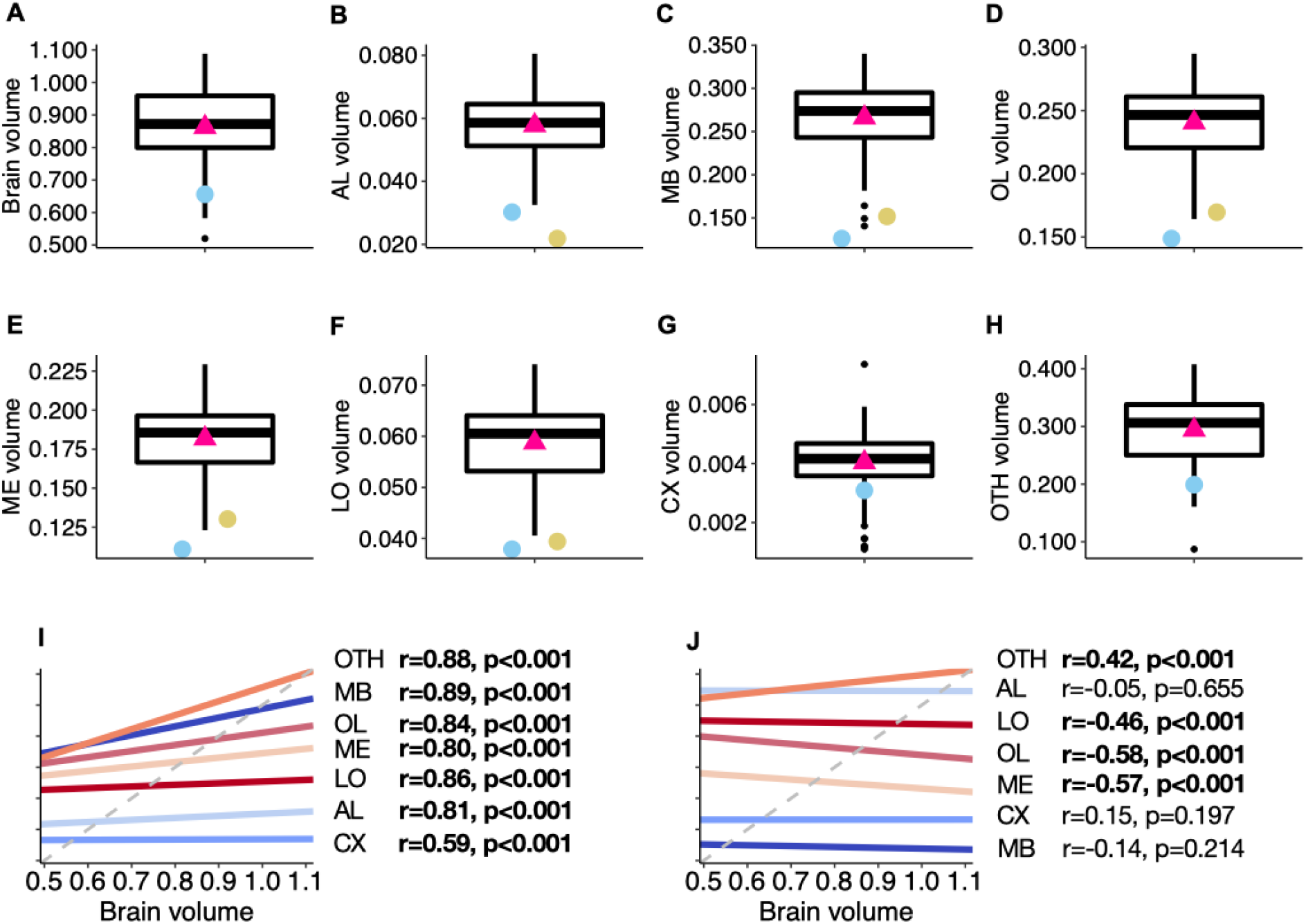
Variation in brain volume and neuropils volumes (mm^3^) and correlation between neuropil volumes and total brain volume for bumblebees (N=77). (**A**) Total brain. (**B**) Antennal lobes (AL). (**C**) Mushroom bodies (MB). (**D**) Optic lobes (OL). (**E**) Medullae (ME). (**F**) Lobulae (LO). (**G**) Central complex (CX). (**H**) Other neuropils (OTH). Boxplots show median volumes (intermediate line) and quartiles (upper and lower lines). Pink triangles display mean volumes. Coloured symbols show the mean of neuropil volumes described for bumblebees in other studies using CT-scan: Rother et al. [48] (N=10 bees - *light blue*), Smith et al. [14] (N=38 bees - *yellow).* (**I**) Linear correlations between absolute volumes of neuropils and total brain volume (mm^3^) (y-axis not given: differs for each neuropil). (**J**) Linear correlations between relative volumes of neuropils and total brain volume (mm^3^) (y-axis not given: differs for each neuropil). The grey dashed line indicates true isometric correlation (slope=1). Pearson correlation coefficient (r) and p-value are given. Strong correlations (r>0.40) and significant correlations (p<0.05) are displayed in bold.

Regarding correlations between neuropil volumes (S5 Table), we found significant and strong positive effects for all possible permutations in absolute volumes. Relative volumes showed the same major trends as in honey bees (S5 Table): OL, ME and LO were positively correlated with each other, OTH was negatively correlated with all other neuropils, except CX, and relative AL volume was positively correlated with MB, OL, ME and LO relative volumes. Thus, the larger the brain, the larger the proportion of OTH and the relatively smaller the other neuropils.

The four bumblebee colonies did not show significant differences for any of the neuropil volumes, except for LO (S4 Table), while we found volume differences for AL, MB, OL and LO when comparing honey bee colonies (Table 1 and Fig 6). Again, this is not surprising since the bumblebee sample was more heterogeneous in terms of individual size than the sample of forager honey bees. Thus, like for honey bees, we found relatively stable intra-colony variability in brain and neuropil volumes (S4 Table).

### Optic lobes showed left-right asymmetry in bumblebees but not in honey bees

Bees have lateralized cognitive abilities, consistently learning better with their right eye [51] and right antenna [52–54]. However, no study has yet reported anatomical asymmetries in the brain that might support these observations [57,59]. Taking advantage of our large number of complete bee brain volumes, we compared the left and right volumes of all paired neuropils in the two bee species (Table 1, Fig 8 and S4 Table). Note that left and right MB structures were often difficult to separate, so the comparison of the two MB sides was only based on 59 honeybees and 36 bumblebees.

**Fig 8.**
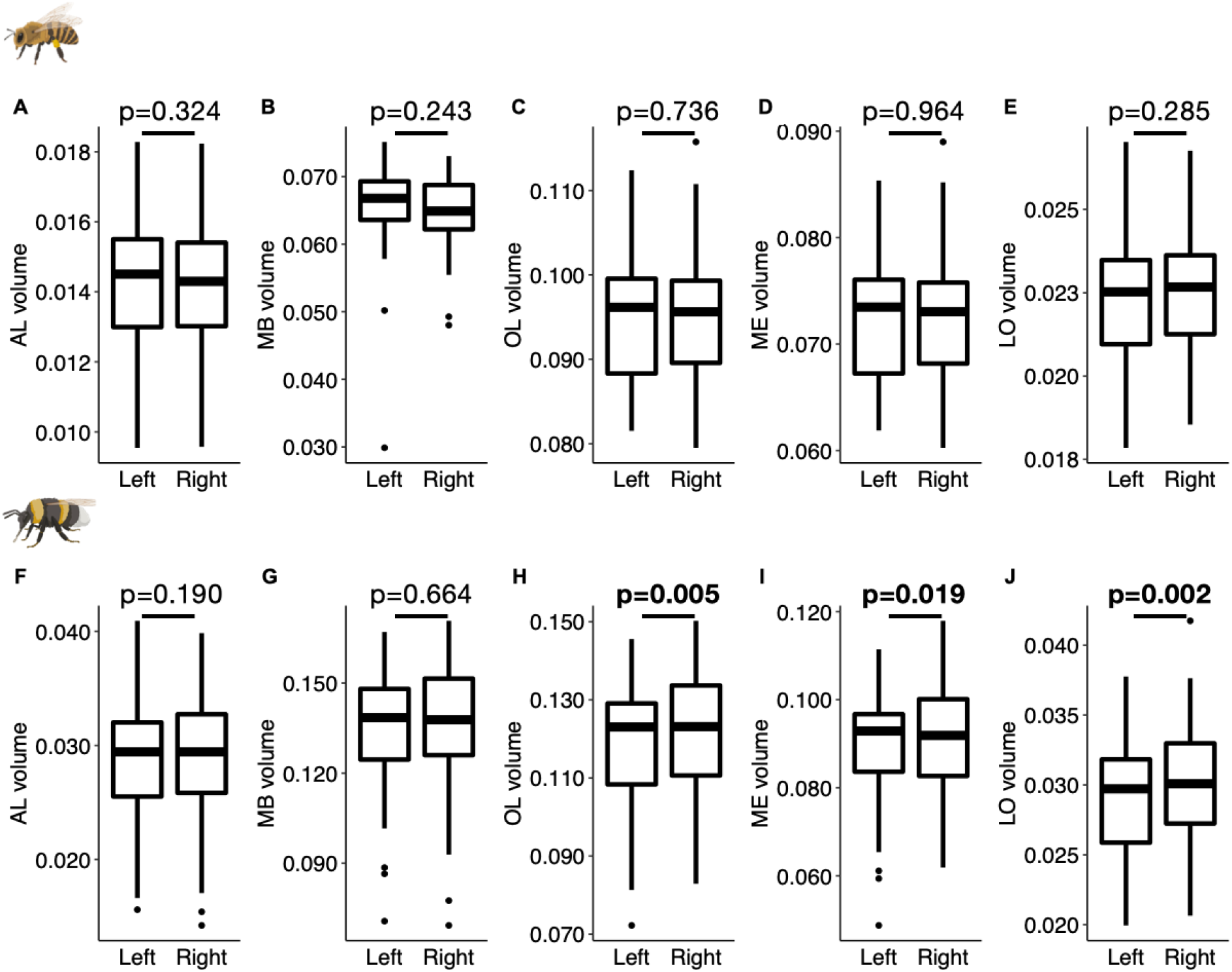
Comparisons between left and right volumes (mm^3^) for paired neuropils in honey bees (N=110; A-E) and bumblebees (N=77; F–J). (**A** and **F**) Antennal lobes (AL). (**B** and **G**) Mushroom bodies (MB). (**C** and **H**) Optic lobes (OL). (**D** and **I**) Medullae (ME). (**E** and **J**) Lobulae (LO). Boxplots show median volumes (intermediate line) and quartiles (upper and lower lines). Statistical differences (p-values) for the neuropil volume between left and right side were obtained with Student’s t-Test, and are displayed in bold when significant.

First, we tested for lateralization of individual brain areas. For honey bee brains, we found no lateral differences in the absolute volumes of AL (LMM: p=0.324), MB (LMM: p=0.243), ME (LMM: p=0.964), LO (LMM: p=0.285) and OL (LMM: p=0.736). We did not find any significant difference in the relative volume of AL (LMM: p=0.341), MB (LMM: p=0.338), ME (LMM: p=0.907), LO (LMM: p=0.245) and OL (LMM: p=0.665) neither. By contrast, the same analysis revealed some asymmetry in bumblebee brains. While there was no difference in the lateral absolute volumes of AL (LMM: p=0.190) and MB (LMM: p=0.664), the left ME (LMM: p=0.019) and left LO were significantly smaller (LMM: p=0.002) in 62% and 66% of the brains, respectively, resulting in an overall smaller left OL (LMM: p=0.005) in 62% of the brains (Fig 8). We found similar results when analysing the relative lateral volumes with no differences in AL (LMM: p=0.247) and MB (LMM: p=0.864), but smaller left ME (LMM: p=0.015) and left LO (LMM: p=0.001), which also results in the left OL having a smaller relative volume than the right side (LMM: p=0.004).

We next investigated potential whole-brain lateralization by comparing the asymmetries along the right-left axis between all paired neuropils (Fig 9). Firstly, we looked for correlations between AL and OL (Figs 9A and 9D). In honey bees, we found an even distribution of individuals across the four potential brain classes: larger right AL and right OL (24.5%), larger right AL and smaller right OL (22.7%), smaller right AL and larger right OL (24.5%), smaller right AL and OL (28.2%). In contrast, we found more bumblebees with larger right AL and larger right OL (39.0%) than in the other categories (S3 Fig). Additionally, we found a positive correlation between the right-left AL and OL volumes (r=0.26, p=0.023; Fig 9D). When taking into account the MB in the analyses, we found no significant correlations between MB and AL or MB and OL in honey bees (Figs 9B and 9C) or bumblebees (Figs 9E and 9F).

**Fig 9.**
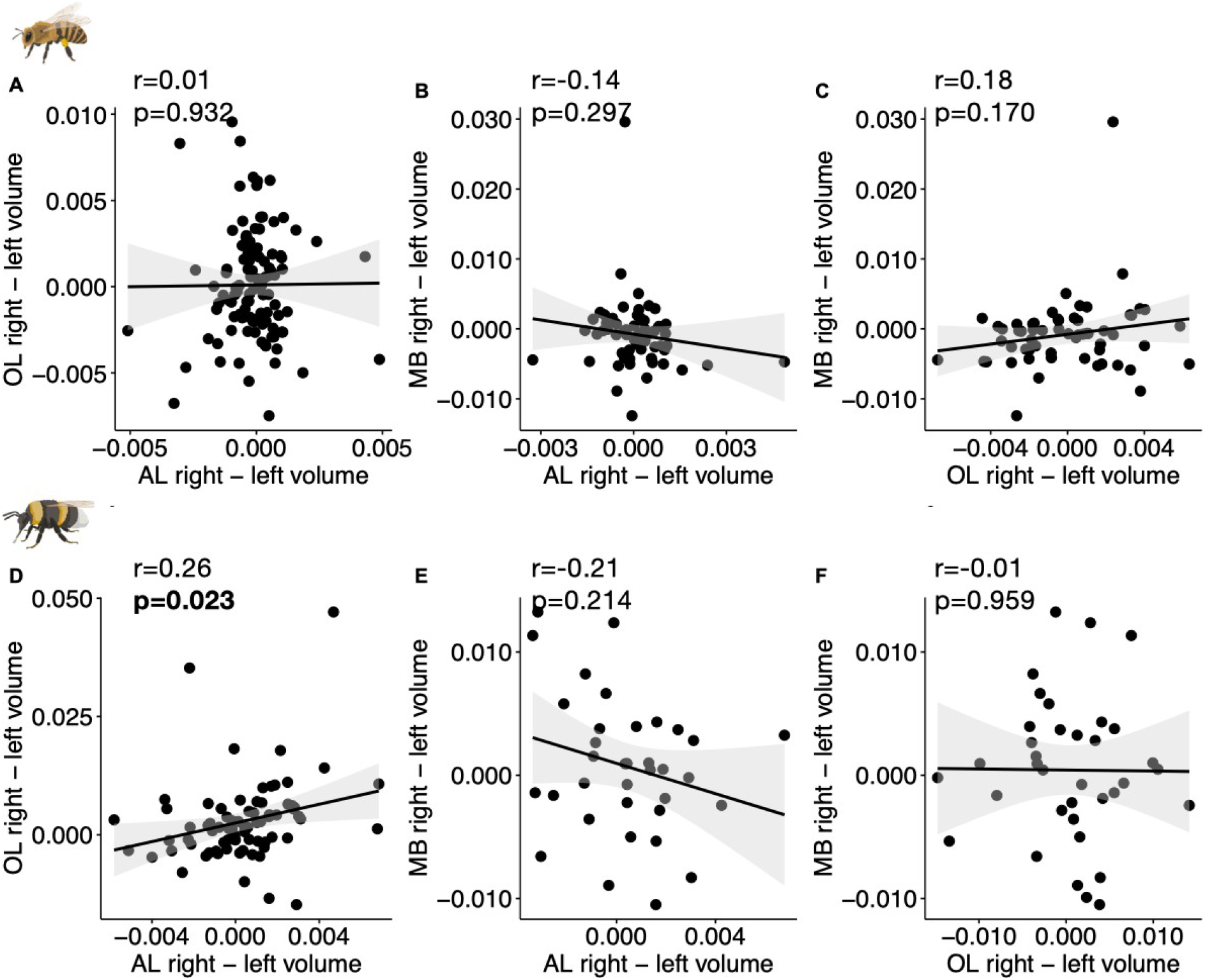
Correlations between right-left volumes (mm^3^) between AL, OL and MB. (**A** and **D**) Correlations between AL right-left and OL right-left volumes (N=110 honey bees, N=77 bumblebees). (**B** and **E**) Correlations between MB right-left and AL right-left volumes (N=59 honey bees, N=36 bumblebees). (**C** and **F**) Correlations between MB right-left and OL right-left volumes (N=59 honey bees, N=36 bumblebees). Pearson correlation coefficient (r) and p-value are given. Strong correlations (r>0.40) and significant correlations (p<0.05) are displayed in bold.

Therefore, there was no clear lateralization at the brain level, except in bumblebees where the majority of individuals had larger right AL and OL. This asymmetry is consistent with inter-individual cognitive variability in bumblebees [55] and the fact that bees have better visual and olfactory learning abilities when stimuli are applied to right sensory organs [51–54]. However, whether this correspondence between neuroanatomy and behaviour is causal remains to be explored [56]. The fact that we did not find such evidence of volumetric asymmetries in honey bees, despite well-known learning asymmetries [51,52], may indicate that left-right differences in ALs or OLs were too small to reach significance in our sample. Alternatively, the reported lateralized learning may be caused by differences between right and left sensory organs (e.g. there are more olfactory sensillae on the right antenna [43]), instead of brain structures.

## Conclusion

Comparative analyses of animal brain volumes are currently limited to a few individuals in model species [57]. Here we show how deep neural networks can be used for fast and accurate analysis of large amounts of volumetric brain images, opening new possibilities for broad comparative analyses in insects and invertebrates in general. Our results on honey bees and bumblebees show high natural variability in brain area sizes that is stable across colonies, suggesting selection for variability supporting adaptive division of labour. Brain asymmetries may also explain previously reported behavioural lateralization [58,59]. Future combination of brain and behavioural analyses using our method for automated image segmentation will help address important open and hotly debated questions related to neuroscience and cognition. Are bigger brains more performant [60]? What are the influences of social and ecological factors in brain evolution [2]? What is the effect of environmental stressors on brain development and cognition [13]? Our image analysis tool, Biomedisa, is accessible via a web browser and does not require complex or tedious software configuration or parameter optimisation, making it user-friendly even for biologists with no substantial computational expertise. We therefore expect our approach to be widely used to study the evolution of animal body parts or organs, beyond the brain itself. Indeed, recent studies showed Biomedisa is suitable for many types of volumetric image data other than micro-CT scans, e.g. from confocal laser scanning microscopy (CLSM), focused ion beam scanning electron microscopy (FIB-SEM), histological imaging [61], or magnetic resonance imaging (MRI) [62], which offer the opportunity for a wide range of studies in biology, ecology and evolution to perform large-scale, cost-effective and time-efficient analyses.

## Materials and Methods

### Sample preparation and CT scan

We performed micro-computed tomography scanning of honey bees (*Apis mellifera*, Buckfast) and bumblebees (*Bombus terrestris*). Honey bees were collected from 9 hives located in two apiaries around Toulouse, France, in August 2020 (Population A (GPS coordinates: 43.55670, 1.47073): 100 bees from 6 hives; population B (43.26756, 2.28376): 20 bees from 3 hives). Foragers returning to the colony were collected at the hive entrance, frozen and stored at −18°C. Bumblebees were purchased from a commercial supplier (Koppert, France) and 77 workers were collected from 4 colonies and frozen at −18°C. For staining bee brains, we removed the front cuticle just above the mandibles and submerged the heads in phosphotungstic acid (5% in a 70/30% ethanol/water solution) for 15 days at ambient temperature [14]. The staining agent is non-hazardous and does not lead to overstaining of the soft tissues, in contrast to other compounds previously used in CT scan studies of insect brains such as osmium-tetroxide [41] or iodine [63]. Two heads were scanned at the same time (as both fit in the field of view of the flat-panel imager) using the micro-CT station EasyTom 150/RX Solutions (Montpellier Ressources Imagerie, France), with the following parameters: resolution of 5.4 μm isotropic voxel size, 40 kV, 130 μA, 736 radiographic projections (acquisition time: 15 minutes). Raw data for each brain scan was reconstructed using X-Act software (RX Solutions, Chavanod, France). The reconstructed volumes were then re-oriented to the same (frontal) plane-of-view and each brain was re-sliced into a new series of two-dimensional images.

### Statistical analysis

We analysed the parameters obtained from the reconstructed neuropils using R Studio v.1.2.5033 [64]. We assessed correlations between brain neuropil volumes using the *rcorr* function from the *Hmisc* package [65]. To analyse the inter-colonial variations of brain volume, we conducted linear mixed models (LMMs) using the *lme4* package [66], with hive as fixed effect and population as random factor. LMMs were followed by F-tests to test the significance of fixed categorical variables using the *anova* function in the *car* package [67]. To assess the potential lateralization of paired neuropils, we conducted Student’s t-Test using Python’s SciPy library.

### Manual processing times with or without Biomedisa

An exclusively manual segmentation of a CT-scan took about 5 to 6 h, the semi-automated segmentation by Biomedisa 2 to 3 h and the manual post-processing of the automatic segmentation results each 5 to 10 min. That means, the manual segmentation of 110 honey bee scans took 550 to 660 h, while semi-automatic creation of three initial training images and the manual post-processing of the automated segmentation results of the remaining 107 CT-scans took at at least 15 h (3 × 2 × 60 min + 107 × 5 min) and a maximum of 27 h (3 × 3 × 60 min + 107 × 10 min). For the segmentation of 77 CT-scans of bumblebee brains, manual segmentation took 385 to 462 h. Here, the creation of 13 training images and the manual post-processing of the remaining 64 automated segmentation results took at least 31 h (13 × 2 × 60 min + 64 × 5 min) and up to 50 h (13 × 3 × 60 min + 64 × 10 min).

### Artificial neural network architecture

Biomedisa uses Keras with TensorFlow backend. A patch-based approach is utilised in which 3D patches of the volumetric images are used instead of the entire CT scan. The patches serve as input for a 3D U-Net and have a size of 64 × 64 × 64 voxels. Before extracting the patches, images are scaled to a size of 256 × 256 × 256 voxels and transformed to have the same mean and variance. An overlapping of the patches is achieved by a stride size of 32 pixels that can be configured in Biomedisa. The network architecture of the deep neural network follows the typical architecture of a 3D U-Net [68]. It consists of a contracting and an expansive part with a repeated application of two 3 × 3 × 3 convolutions, each followed by batch normalisation and a rectified linear unit (ReLU) activation layer. Each contracting block is followed by a 2 × 2 × 2 max pooling operation with stride size of 2 for downsampling. At each downsampling step, the number of feature channels is doubled, starting with 32 channels. Every step in the expansive part consists of an upsampling of the feature map and a concatenation with the corresponding cropped feature map from the contracting path, followed by two 3 × 3 × 3 convolutions, with each followed by batch normalisation and a ReLU activation layer. At the final layer, a 1 × 1 × 1 convolution is used to map each feature vector to the desired number of labels. To train the network, stochastic gradient descent is used with a learning rate of 0.01, decay of 1 × 10^-6^, momentum of 0.9, enabled Nesterov momentum, 200 training epochs, and a batch size of 24 samples.

### Removing outliers

To evaluate the segmentation results, Biomedisa’s cleanup function was used for all neuropils except CX to automatically remove outliers or islands with a threshold of 0.1. This threshold value was set lower than the standard configuration (0.9) in order to avoid a partial deletion of paired neuropils.

### Evaluating the creation of training data

To evaluate the semi-automatic segmentation used to create the labels for the training images, we compared the commonly used linear interpolation in AVIZO 2019.1 and the smart interpolation in Biomedisa using the set of 26 training images. For intervals including CX we used every 5th slice of the ground truth data, otherwise every 10th slice as initialisation. Using the same pre-segmented slices, Biomedisa’s smart interpolation, which also takes into account the underlying image data, achieves higher segmentation accuracy (average Dice score of 0.967) compared to the purely morphological interpolation of AVIZO 2019.1 (average Dice score of 0.928, Fig 3 and S1 Table). Thus, the manual work required to create the training data for a neural network is significantly reduced.

### Increasing the number of training images

To test the performance of the automatic segmentation in terms of the number of training images, we evaluated the accuracy of the trained network using 3 to 26 training images (Fig 3 and S1 Table). Here, the 110 honey bee images and the corresponding labels were split into 26 training images, 30 validation images and 54 test images. The accuracy of the training process, measured using the Dice score, was evaluated using the validation data after each of the 200 epochs performed. Only the best performing network was saved. Finally, the trained network was applied to the 54 test images and the average Dice score of the segmentation results was calculated. While increasing the number of training images improves the accuracy of the automatic segmentation and thus reduces the required manual post-processing, this improvement gradually slows down with an increasing number of training images. Therefore, one has to weigh the gain in accuracy against the additional effort required. Using the honey bee dataset, 12 to 20 training images are sufficient for adequate automatic segmentation.

### Uncropped image data

The network was also trained and evaluated using uncropped images (i.e. original honey bee CT scans). Cropping the image data to the area of the neuropils significantly increased the segmentation accuracy of the neural network from a total Dice score of 0.928 for the uncropped images to 0.970 for the cropped image data (Fig 3 and S1 Table). By default, Biomedisa scales each image to a size of 256 pixels for each axis to facilitate training. The cropped image data (average size of 451 × 273 × 167 voxels) were thus scaled with an averaged factor of 0.57 and 1.53 along the x- and z-axes, and only marginally along the y-axis. Without cropping the image data (average size of 844 × 726 × 485 voxels), a large amount of redundant information is added to the training data. In addition, the loss of resolution due to the scaling of the image is significantly larger compared to cropped image data.

### Automatic cropping

As an alternative to manual cropping, a neural network can be trained with Biomedisa to automatically crop the image data to the region of interest before segmentation. Here the DenseNet121 [69] pre-trained on the ImageNet database is used to decide whether a slice of the volume contains the object or not. Applying the network to all three axes creates a bounding box covering the honey bee brain. Adding a buffer of 25 slices to all sides after cropping ensures that the entire object is within the bounding box. The network is trained in two steps. First, the head of the pre-trained network is removed and replaced with a global average pooling layer, a dropout layer with a probability of 0.3 to avoid overfitting, and a final dense layer with sigmoid activation function. The weights of the base model are kept fixed while the head is optimised with binary cross-entropy, Adam optimiser with a learning rate of 0.001 and 50 epochs. Second, the entire DenseNet121 is fine-tuned to the honey bee training data with a learning rate of 1 × 10^-5^ and 50 epochs. Auto-cropping achieves a Dice score of 0.952, which increases accuracy by 2.4% compared to uncropped honey bee image data (Fig 3 and S1 Table).

### Evaluation metrics

For two segmentations *X* and *χ′* consisting of *n* labels, the Dice similarity coefficient (Dice) is defined as

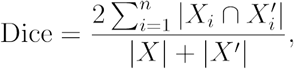

where |*X*| and |*χ′*| are the total number of voxels of each segmentation, respectively, and *X*_i_, is the subset of voxels of *X* with label *i*. For the surfaces *S*_i_, and 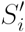 of the two segmentations, the average symmetric surface distance (ASSD) is defined as

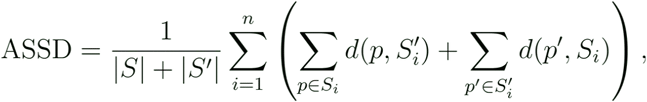

where

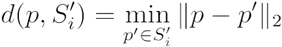

is the Euclidean distance from a point *p* on the surface *S*_i_, of label *i* to the closest point *p*′ on the corresponding surface 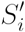 of the second segmentation.

### Denoising image data

Volumetric images are denoised using an arithmetic mean filter

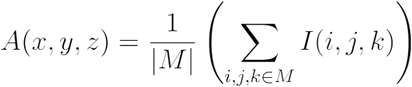

with a filter mask *M* of 3 × 3 × 3 voxels.

## Data Availability Statement

Image and label data of honey bees and bumblebees as well as trained networks are available at https://biomedisa.org/gallery/. The source code is available as part of the Biomedisa open source project. It was developed and tested for Ubuntu 20.04 LTS and Windows 10. Biomedisa can be used via the command line or with any common browser as an interface. The source code can be downloaded from https://github.com/biomedisa/biomedisa/ and installed according to the installation instructions. All other relevant data are within the paper and its Supporting Information files.

## Abbreviations

AL: antennal lobes
MB: mushroom bodies
CX: central complex
ME: medullae
LO: lobulae
OL: optic lobes
OTH: other neuropils

## Author contributions

Conceptualization: Philipp D. Lösel, Coline Monchanin, Renaud Lebrun, Jean-Marc Devaud, Vincent Heuveline, Mathieu Lihoreau.

Data curation: Philipp D. Lösel, Coline Monchanin.

Formal analysis: Philipp D. Lösel, Coline Monchanin.

Funding acquisition: Philipp D. Lösel, Vincent Heuveline, Mathieu Lihoreau.

Investigation: Philipp D. Lösel, Coline Monchanin.

Methodology: Philipp D. Lösel, Coline Monchanin.

Project Administration: Philipp D. Lösel, Vincent Heuveline, Mathieu Lihoreau.

Resources: Philipp D. Lösel, Coline Monchanin, Vincent Heuveline.

Software: Philipp D. Lösel, Coline Monchanin, Alejandra Jayme, Jacob Relle.

Supervision: Vincent Heuveline, Mathieu Lihoreau.

Validation: Philipp D. Lösel.

Visualization: Philipp D. Lösel, Coline Monchanin.

Writing – original draft: Philipp D. Lösel, Coline Monchanin.

Writing – review & editing: Philipp D. Lösel, Coline Monchanin, Renaud Lebrun, Alejandra Jayme, Jean-Marc Devaud, Mathieu Lihoreau.

## Materials & Correspondence

Correspondence and requests for materials should be addressed to P.D.L. or C.M.

## References

1. Appenzeller T. The AI revolution in science [Internet]. 2017 [cited December 2022 Dec 23]. Available from: https://doi.org/10.1126/science.aan7064.

2. Dunbar RIM. The social brain hypothesis. Evol Anthropol Issues News Rev. 1998; 6(5):178–90. https://doi.org/10.1002/(SICI)1520-6505(1998)6:5<178::AID-EVAN5>3.0.CO;2-8

3. Cardinale B. Impacts of biodiversity loss. Science. 2012; 336(6081):552–3. https://doi.org/10.1126/science.1222102

4. Kumazawa-Manita N, Katayama M, Hashikawa T, Iriki A. Three-dimensional reconstruction of brain structures of the rodent *Octodon degus:* a brain atlas constructed by combining histological and magnetic resonance images. Exp Brain Res. 2013; 231(1):65–74. https://doi.org/10.1007/s00221-013-3667-1

5. Newman JD, Kenkel WM, Aronoff EC, Bock NA, Zametkin MR, Silva AC. A combined histological and MRI brain atlas of the common marmoset monkey, *Callithrix jacchus*. Brain Res Rev. 2009; 62(1):1–18. https://doi.org/10.1016/j.brainresrev.2009.09.001

6. Rein K, Zöckler M, Mader MT, Grübel C, Heisenberg M. The *Drosophila* standard brain. Curr Biol. 2002; 12(3):227–31. https://doi.org/10.1016/s0960-9822(02)00656-5

7. Healy SD. Adaptation and the brain. 1st ed. Oxford University Press; 2021. https://doi.org/10.1093/oso/9780199546756.001.0001

8. dos Santos Rolo T, Ershov A, van de Kamp T, Baumbach T. In vivo X-ray cine-tomography for tracking morphological dynamics. Proc Natl Acad Sci. 2014; 111(11):3921–6. https://doi.org/10.1073/pnas.1308650111

9. Maire E, Withers PJ. Quantitative X-ray tomography. Int Mater Rev. 2014; 59(1):1–43. https://doi.org/10.1179/1743280413Y.0000000023

10. Richter A, Hita Garcia F, Keller RA, Billen J, Economo EP, Beutel RG. Comparative analysis of worker head anatomy of Formica and Brachyponera (Hymenoptera: Formicidae). Arthropod Syst Phylogeny. 2020; 78(1):133–70. http://www.doi.org/10.26049/ASP78-1-2020-06

11. van de Kamp T, Schwermann AH, dos Santos Rolo T, Lösel PD, Engler T, Etter W, et al. Parasitoid biology preserved in mineralized fossils. Nat Commun. 2018; 9(1):3325. https://doi.org/10.1038/s41467-018-05654-y

12. van de Kamp T, Cecilia A, dos Santos Rolo T, Vagovič P, Baumbach T, Riedel A. Comparative thorax morphology of death-feigning flightless cryptorhynchine weevils (Coleoptera: Curculionidae) based on 3D reconstructions. Arthropod Struct Dev. 2015; 44(6):509–23. https://doi.org/10.1016/j.asd.2015.07.004

13. Smith DB, Arce AN, Ramos Rodrigues A, Bischoff PH, Burris D, Ahmed F, et al. Insecticide exposure during brood or early-adult development reduces brain growth and impairs adult learning in bumblebees. Proc R Soc B Biol Sci. 2020; 287(1922):20192442. https://doi.org/10.1098/rspb.2019.2442

14. Smith DB, Bernhardt G, Raine NE, Abel RL, Sykes D, Ahmed F, et al. Exploring miniature insect brains using micro-CT scanning techniques. Sci Rep. 2016; 6(1):21768. https://doi.org/10.1038/srep21768

15. Dumbravă MD, Rothschild BM, Weishampel DB, Csiki-Sava Z, Andrei RA, Acheson KA, et al. A dinosaurian facial deformity and the first occurrence of ameloblastoma in the fossil record. Sci Rep. 2016; 6(1):29271. https://doi.org/10.1038/srep29271

16. Gross V, Müller M, Hehn L, Ferstl S, Allner S, Dierolf M, et al. X-ray imaging of a water bear offers a new look at tardigrade internal anatomy. Zool Lett. 2019; 5(1):14. https://doi.org/10.1186/s40851-019-0130-6

17. Jones MEH, Button DJ, Barrett PM, Porro LB. Digital dissection of the head of the rock dove (Columba livia) using contrast-enhanced computed tomography. Zool Lett. 2019; 5(1):17. https://doi.org/10.1186/s40851-019-0129-z

18. Pardo JD, Szostakiwskyj M, Ahlberg PE, Anderson JS. Hidden morphological diversity among early tetrapods. Nature. 2017; 546(7660):642–5. https://doi.org/10.1038/nature22966

19. Lösel P, Heuveline V. Enhancing a diffusion algorithm for 4D image segmentation using local information. Proc SPIE. 2016; 9784(1):97842L. https://doi.org/10.1117/12.2216202

20. Lösel PD. GPU-basierte Verfahren zur Segmentierung biomedizinischer Bilddaten [Dissertation]. Heidelberg: Heidelberg University; 2022. https://doi.org/10.11588/heidok.00031525

21. Lösel PD, van de Kamp T, Jayme A, Ershov A, Faragó T, Pichler O, et al. Introducing Biomedisa as an open-source online platform for biomedical image segmentation. Nat Commun. 2020; 11(1):5577. https://doi.org/10.1038/s41467-020-19303-w

22. Jandt JM, Bengston S, Pinter-Wollman N, Pruitt JN, Raine NE, Dornhaus A, et al. Behavioural syndromes and social insects: personality at multiple levels. Biol Rev. 2014; 89(1):48–67. https://doi.org/10.1111/brv.12042

23. Fahrbach SE, Moore D, Capaldi EA, Farris SM, Robinson GE. Experience-expectant plasticity in the mushroom bodies of the honeybee. Learn Mem. 1998; 5(1):115–23. PMID: http://www.ncbi.nlm.nih.gov/pubmed/10454376

24. Withers GS, Fahrbach SE, Robinson GE. Selective neuroanatomical plasticity and division of labour in the honeybee. Nature. 1993; 364(6434):238–40. https://doi.org/10.1038/364238a0

25. Giurfa M. Cognition with few neurons: higher-order learning in insects. Trends Neurosci. 2013; 36(5):285–94. https://doi.org/10.1016/j.tins.2012.12.011

26. Menzel R. The honeybee as a model for understanding the basis of cognition. Nat Rev Neurosci. 2012; 13(11):758–68. https://doi.org/10.1038/nrn3357

27. Sombke A, Lipke E, Michalik P, Uhl G, Harzsch S. Potential and limitations of X-Ray micro-computed tomography in arthropod neuroanatomy: a methodological and comparative survey. J Comp Neurol. 2015; 523(8):1281–95. https://doi.org/10.1002/cne.23741

28. Gowda V, Gronenberg W. Brain composition and scaling in social bee species differing in body size. Apidologie. 2019; 50(6):779–92. https://doi.org/10.1007/s13592-019-00685-w

29. Gronenberg W, Couvillon MJ. Brain composition and olfactory learning in honey bees. Neurobiol Learn Mem. 2010; 93(3):435–43. https://doi.org/10.1016/j.nlm.2010.01.001

30. Brandt R, Rohlfing T, Rybak J, Krofczik S, Maye A, Westerhoff M, et al. Three-dimensional average-shape atlas of the honeybee brain and its applications. J Comp Neurol. 2005; 492(1):1–19. https://doi.org/10.1002/cne.20644

31. Strausfeld NJ. Arthropod brains: evolution, functional elegance, and historical significance. Cambridge, Mass: Harvard University Press; 2012. https://doi.org/10.2307/j.ctv1dp0v2h

32. Maier-Hein L, Eisenmann M, Reinke A, Onogur S, Stankovic M, Scholz P, et al. Why rankings of biomedical image analysis competitions should be interpreted with care. Nat Commun. 2018; 9(1):5217. https://doi.org/10.1038/s41467-018-07619-7

33. Robinson GE. Regulation of honey bee age polyethism by juvenile hormone. Behav Ecol Sociobiol. 1987; 20(5):329–38. https://doi.org/10.1007/BF00300679

34. Waddington KD. Implications of variation in worker body size for the honey bee recruitment system. J Insect Behav. 1989; 2(1):91–103. https://doi.org/10.1007/BF01053620

35. Haddad D, Schaupp F, Brandt R, Manz G, Menzel R, Haase A. NMR imaging of the honeybee brain. J Insect Sci. 2004; 4(1):1–7. https://doi.org/10.1093/jis/4.1.7

36. Steijven K, Spaethe J, Steffan-Dewenter I, Härtel S. Learning performance and brain structure of artificially-reared honey bees fed with different quantities of food. PeerJ. 2017; 5:e3858. https://doi.org/10.7717/peerj.3858

37. Mares S, Ash L, Gronenberg W. Brain allometry in bumblebee and honey bee workers. Brain Behav Evol. 2005; 66(1):50–61. https://doi.org/10.1159/000085047

38. Maleszka J, Barron AB, Helliwell PG, Maleszka R. Effect of age, behaviour and social environment on honey bee brain plasticity. J Comp Physiol A. 2009; 195(8):733–40. https://doi.org/10.1007/s00359-009-0449-0

39. Durst C, Eichmüller S, Menzel R. Development and experience lead to increased volume of subcompartments of the honeybee mushroom body. Behav Neural Biol. 1994; 62(3):259–63. https://doi.org/10.1016/s0163-1047(05)80025-1

40. Greco MK, Stait-Gardner T. Applying x-ray micro-tomography to learning and memory. Biomed Phys Eng Express. 2017; 3(2):024001. https://doi.org/10.1088/2057-1976/aa6307

41. Ribi W, Senden TJ, Sakellariou A, Limaye A, Zhang S. Imaging honey bee brain anatomy with micro-X-ray-computed tomography. J Neurosci Methods. 2008; 171(1):93–7. https://doi.org/10.1016/j.jneumeth.2008.02.010

42. Prakash YS, Smithson KG, Sieck GC. Application of the Cavalieri principle in volume estimation using laser confocal microscopy. NeuroImage. 1994 Nov; 1(4):325–33. https://doi.org/10.1006/nimg.1994.1017

43. Barrett M, Schneider S, Sachdeva P, Gomez A, Buchmann S, O’Donnell S. Neuroanatomical differentiation associated with alternative reproductive tactics in male arid land bees, *Centris pallida* and *Amegilla dawsoni*. J Comp Physiol A. 2021; 207(4):497–504. https://doi.org/10.1007/s00359-021-01492-4

44. Nowogrodzki R. Division of labour in the honeybee colony: a review. Bee World. 1984; 65(3):109–16. https://doi.org/10.1080/0005772X.1984.11098788

45. Ismail N, Robinson GE, Fahrbach SE. Stimulation of muscarinic receptors mimics experience-dependent plasticity in the honey bee brain. Proc Natl Acad Sci. 2006; 103(1):207–11. https://doi.org/10.1073/pnas.0508318102

46. Winnington AP, Napper RM, Mercer AR. Structural plasticity of identified glomeruli in the antennal lobes of the adult worker honey bee. J Comp Neurol. 1996; 365(3):479–90. https://doi.org/10.1002/(sici)1096-9861(19960212)365:3%3C479::aid-cne10%3E3.0.co;2-m

47. Goulson D. Bumblebees: behaviour, ecology, and conservation. 2nd ed. Oxford; New York: Oxford University Press; 2010.

48. Rother L, Kraft N, Smith DB, el Jundi B, Gill RJ, Pfeiffer K. A micro-CT-based standard brain atlas of the bumblebee. Cell Tissue Res. 2021; 386(1):29–45. https://doi.org/10.1007/s00441-021-03482-z

49. Austin MW, Dunlap AS. Intraspecific variation in worker body size makes North American bumble bees *(Bombus* spp.) less susceptible to decline. Am Nat. 2019; 194(3):381–94. https://doi.org/10.1086/704280

50. Spaethe J, Weidenmüller A. Size variation and foraging rate in bumblebees (*Bombus terrestris)*. Insectes Sociaux. 2002; 49(2):142–6. https://doi.org/10.1007/s00040-002-8293-z

51. Letzkus P, Boeddeker N, Wood JT, Zhang SW, Srinivasan MV. Lateralization of visual learning in the honeybee. Biol Lett. 2008; 4(1):16–9. https://doi.org/10.1098/rsbl.2007.0466

52. Letzkus P, Ribi WA, Wood JT, Zhu H, Zhang SW, Srinivasan MV. Lateralization of olfaction in the honeybee *Apis mellifera*. Curr Biol. 2006; 16(14):1471–6. https://doi.org/10.1016/j.cub.2006.05.060

53. Anfora G, Rigosi E, Frasnelli E, Ruga V, Trona F, Vallortigara G. Lateralization in the invertebrate brain: left-right asymmetry of olfaction in bumble bee, *Bombus terrestris*. Burne T, editor. PLoS ONE. 2011; 6(4):e18903. https://doi.org/10.1371/journal.pone.0018903

54. Rogers LJ, Vallortigara G. From antenna to antenna: lateral shift of olfactory memory recall by honeybees. PLoS ONE. 2008; 3(6):e2340. https://doi.org/10.1371/journal.pone.0002340

55. Chittka L, Dyer AG, Bock F, Dornhaus A. Bees trade off foraging speed for accuracy. Nature. 2003; 424(6947):388–388. https://doi.org/10.1038/424388a

56. Smith KE, Raine NE. A comparison of visual and olfactory learning performance in the bumblebee *Bombus terrestris*. Behav Ecol Sociobiol. 2014; 68(9):1549–59. https://doi.org/10.1007/s00265-014-1765-0

57. Sahin B, Aslan H, Unal B, Canan S, Bilgic S, Kaplan S, et al. Brain volumes of the lamb, rat and bird do not show hemispheric asymmetry: a stereological study. Image Anal Stereol. 2011; 20(1):9. https://doi.org/10.5566/ias.v20.p9-13

58. Frasnelli E, Vallortigara G, Rogers LJ. Left–right asymmetries of behaviour and nervous system in invertebrates. Neurosci Biobehav Rev. 2012; 36(4):1273–91. https://doi.org/10.1016/j.neubiorev.2012.02.006

59. Frasnelli E, Haase A, Rigosi E, Anfora G, Rogers L, Vallortigara G. The bee as a model to investigate brain and behavioural asymmetries. Insects. 2014; 5(1):120–38. https://doi.org/10.3390/insects5010120

60. Chittka L, Niven J. Are bigger brains better? Curr Biol. 2009; 19(21):R995–1008. https://doi.org/10.1016/j.cub.2009.08.023

61. Csader M, Mayer K, Betz O, Fischer S, Eggs B. Ovipositor of the braconid wasp Habrobracon hebetor: structural and functional aspects. J Hymenopt Res. 2021; 83:73–99. https://doi.org/10.3897/jhr.83.64018

62. Lösel P, Heuveline V. A GPU Based Diffusion Method for Whole-Heart and Great Vessel Segmentation. In: Zuluaga MA, Bhatia K, Kainz B, Moghari MH, Pace DF, editors. Reconstruction, Segmentation, and Analysis of Medical Images. Cham: Springer International Publishing; 2017. p. 121–8. https://doi.org/10.1007/978-3-319-52280-7_12

63. Lesciotto KM, Motch Perrine SM, Kawasaki M, Stecko T, Ryan TM, Kawasaki K, et al. Phosphotungstic acid-enhanced microCT: optimized protocols for embryonic and early postnatal mice. Dev Dyn. 2020; 249(4):573–85. https://doi.org/10.1002/dvdy.136

64. RStudio Team. RStudio: integrated development for R. Boston, MA: RStudio, PBC; 2020. Available from: http://www.rstudio.com/.

65. Harrell FE. Hmisc: Harrell Miscellaneous [Internet]. 2022 [cited 2022 Dec 21]. Available from: https://cran.r-project.org/web/packages/Hmisc/Hmisc.pdf.

66. Bates D, Mächler M, Bolker B, Walker S. Fitting linear mixed-effects models using lme4. J Stat Softw. 2015; 67(1):1–48. https://doi.org/10.18637/jss.v067.i01

67. Fox J, Weisberg S. An R companion to applied regression. 3rd ed. Los Angeles: SAGE; 2019.

68. Ronneberger O, Fischer P, Brox T. U-Net: Convolutional Networks for Biomedical Image Segmentation. In: Navab N, Hornegger J, Wells WM, Frangi AF, editors. Medical Image Computing and Computer-Assisted Intervention – MICCAI 2015. Cham: Springer International Publishing; 2015. p. 234–41. https://doi.org/10.1007/978-3-319-24574-4_28

69. Huang G, Liu Z, Van Der Maaten L, Weinberger KQ. Densely connected convolutional networks. In: 2017 IEEE Conference on Computer Vision and Pattern Recognition (CVPR). Honolulu, HI: IEEE; 2017. p. 2261–9. https://doi.org/10.1109/CVPR.2017.243

