## Supplementary material for "Natural variability in bee brain size and symmetry revealed by micro-CT imaging and deep learning": S1 Table

### Supporting Information

**S1 Table. Average Dice scores of semi-automatic and automatic segmentation results of bumblebee (row 2) and honey bee brains (rows 3–12).** Outliers were automatically removed (see "Methods"). Last column: amount of manual correction required (Error in % of Dice score). Brain areas are labelled using the same abbreviations as in Fig 2. For some of the performance tests of the automatic segmentation (*yellow*), the 84 honey bee test images were split into 30 validation images and 54 test images (see "Methods").

| Dataset | AL | MB | ME | LO | CX | OTH | Total | Error |
|---|---|---|---|---|---|---|---|---|
| **64 test images** 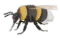 | 0.983 | 0.985 | 0.983 | 0.982 | 0.837 | 0.980 | **0.983** | **1.7%** |
| **84 test images** 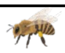 | 0.984 | 0.982 | 0.992 | 0.991 | 0.967 | 0.988 | **0.988** | **1.2%** |
| **3 training images** | 0.841 | 0.890 | 0.954 | 0.940 | 0.746 | 0.917 | **0.919** | **8.1%** |
| **7 training images** | 0.946 | 0.926 | 0.973 | 0.965 | 0.816 | 0.950 | **0.951** | **4.9%** |
| **12 training images** | 0.947 | 0.942 | 0.975 | 0.977 | 0.882 | 0.958 | **0.960** | **4.0%** |
| **18 training images** | 0.960 | 0.947 | 0.979 | 0.981 | 0.875 | 0.963 | **0.965** | **3.5%** |
| **26 training images** | 0.968 | 0.953 | 0.982 | 0.984 | 0.906 | 0.968 | **0.970** | **3.0%** |
| **Uncropped data** | 0.924 | 0.898 | 0.956 | 0.941 | 0.775 | 0.924 | **0.928** | **7.2%** |
| **Automatic cropping** | 0.949 | 0.928 | 0.970 | 0.968 | 0.865 | 0.951 | **0.952** | **4.8%** |
| **Biomedisa interpolation** | 0.967 | 0.949 | 0.986 | 0.982 | 0.856 | 0.962 | **0.967** | **3.3%** |
| **AVIZO interpolation** | 0.925 | 0.925 | 0.915 | 0.914 | 0.848 | 0.946 | **0.928** | **7.2%** |
