## Supplementary material for "Natural variability in bee brain size and symmetry revealed by micro-CT imaging and deep learning": S2 Table

**S2 Table. Correlation between absolute neuropil volumes (bottom left) and between relative neuropil volumes (top right) for honey bees (N=110).** Pearson correlation coefficient and p-value are given. Strong correlations (r>0.40) and significant correlations (p<0.05) are displayed in bold. Brain areas are labelled using the same abbreviations as in Fig 2.

| 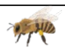 | AL | MB | OL | ME | LO | CX | OTH |
|---|---|---|---|---|---|---|---|
| **AL** | | **-0.45** **(p<0.001)** | 0.27 **(p=0.005)** | 0.24 **(p=0.010)** | 0.27 **(p=0.004)** | 0.08 (p=0.424) | -0.06 (p=0.511) |
| **MB** | 0.22 **(p=0.019)** | | -0.26 **(p=0.006)** | -0.27 **(p=0.005)** | -0.18 (p=0.061) | -0.01 (p=0.901) | **-0.54** **(p<0.001)** |
| **OL** | **0.71** **(p<0.001)** | 0.35 **(p<0.001)** | | **0.98** **(p<0.001)** | **0.81** **(p<0.001)** | 0.21 **(p=0.025)** | **-0.65** **(p<0.001)** |
| **ME** | **0.69** **(p<0.001)** | 0.33 **(p<0.001)** | **0.99** **(p<0.001)** | | **0.69** **(p<0.001)** | 0.16 (p=0.088) | **-0.62** **(p<0.001)** |
| **LO** | **0.69** **(p<0.001)** | 0.39 **(p<0.001)** | **0.93** **(p<0.001)** | **0.88** **(p<0.001)** | | 0.32 **(p<0.001)** | **-0.57** **(p<0.001)** |
| **CX** | 0.32 **(p<0.001)** | 0.13 (p=0.168) | 0.35 **(p<0.001)** | 0.32 **(p<0.001)** | **0.43** **(p<0.001)** | | -0.21 **(p=0.025)** |
| **OTH** | **0.60** **(p<0.001)** | 0.38 **(p<0.001)** | **0.59** **(p<0.001)** | **0.57** **(p<0.001)** | **0.57** **(p<0.001)** | 0.30 **(p=0.002)** | |
