## Supplementary material for "Natural variability in bee brain size and symmetry revealed by micro-CT imaging and deep learning": S3 Table

**S3 Table. Inter-individual variability of total brain and neuropil volumes (%) within honey bee hives.** Percentage of volume variation ((Max-Min)*100/Max) was calculated for total brain and all neuropils for each hive. Brain areas are labelled using the same abbreviations as in Fig 2.

| Population | Hive | Number of bees | Brain volume | AL | MB | OL | CX | OTH |
|---|---|---|---|---|---|---|---|---|
| A | H1 | 9 | 11.52% | 37.23% | 14.01% | 16.90% | 37.87% | 17.45% |
| A | H2 | 11 | 26.60% | 33.34% | 30.22% | 14.28% | 19.75% | 39.72% |
| A | H3 | 18 | 26.32% | 20.51% | 22.09% | 17.36% | 27.48% | 50.61% |
| A | H4 | 19 | 27.02% | 28.36% | 21.20% | 25.21% | 49.36% | 36.37% |
| A | H5 | 17 | 20.66% | 31.38% | 27.23% | 21.44% | 31.17% | 31.56% |
| A | H6 | 16 | 17.18% | 33.70% | 26.11% | 14.08% | 39.97% | 21.82% |
| B | H7 | 6 | 18.33% | 33.60% | 15.27% | 17.98% | 25.96% | 19.74% |
| B | H8 | 11 | 23.82% | 22.68% | 29.70% | 19.61% | 26.45% | 30.70% |
| B | H9 | 3 | 8.25% | 15.00% | 6.70% | 9.09% | 11.30% | 11.02% |
