## Supplementary material for "Natural variability in bee brain size and symmetry revealed by micro-CT imaging and deep learning": S4 Table

**S4 Table. Total brain and neuropil volumes of bumblebees (N=77).** Mean (± standard deviation), minimal and maximal volumes ($mm^3$). Global percentage of volume variation and inter-individual variability of brain and neuropils volume (%) within colonies (N=77 bumblebees). Information is also given for the left and right sides of paired neuropils (i.e. AL, MB, OL, ME, LO). F-test, following LMMs, tests the significance of the fixed variable 'colony', and results are displayed in bold when significant. Brain areas are labelled using the same abbreviations as in Fig 2.

| **Volume ($mm^3$)** | **Mean ± s.d.** | **Min** | **Max** | **% variation** | **F-test** | **Colony A (N=18)** | **Colony B (N=21)** | **Colony C (N=21)** | **Colony D (N=18)** |
|---|---|---|---|---|---|---|---|---|---|
| **Brain** | 0.862±0.127 | 0.520 | 1.089 | 52.27% | F(3,73)=1.854, p=0.145 | 42.53% | 43.08% | 51.65% | 29.79% |
| **AL** | 0.058±0.010 | 0.033 | 0.081 | 59.63% | F(3,73)=1.492, p=0.224 | 55.30% | 55.01% | 57.65% | 53.25% |
| ***Left*** | *0.029±0.005* | *0.016* | *0.041* | *61.90%* | | | | | |
| ***Right*** | *0.029±0.006* | *0.014* | *0.040* | *64.33%* | | | | | |
| **MB** | 0.266±0.041 | 0.141 | 0.340 | 58.70% | F(3,73)=2.004, p=0.121 | 49.47% | 41.35% | 58.70% | 36.19% |
| ***Left*** | *0.135±0.021* | *0.071* | *0.167* | *57.70%* | | | | | |
| ***Right*** | *0.135±0.023* | *0.069* | *0.171* | *59.44%* | | | | | |
| **OL** | 0.240±0.030 | 0.164 | 0.295 | 44.34% | F(3,73)=2.340, p=0.080 | 30.57% | 43.90% | 38.63% | 31.28% |
| ***Left*** | *0.119±0.016* | *0.072* | *0.146* | *50.38%* | | | | | |
| ***Right*** | *0.122±0.015* | *0.082* | *0.150* | *44.83%* | | | | | |
| **ME** | 0.182±0.023 | 0.123 | 0.229 | 46.37% | F(3,73)=2.014, p=0.119 | 32.26% | 46.37% | 37.47% | 31.38% |
| ***Left*** | *0.090±0.012* | *0.049* | *0.111* | *56.27%* | | | | | |
| ***Right*** | *0.092±0.011* | *0.06* | *0.118* | *47.52%* | | | | | |

103
104

| | | | | | | | | | |
|---|---|---|---|---|---|---|---|---|---|
| | | *2* | | | | | | | |
| **LO** | 0.059±0.008 | 0.040 | 0.074 | 45.25% | F(3,73)=2.750, p=0.049 | 37.10% | 44.40% | 44.41% | 31.81% |
| ***Left*** | *0.029±0.004* | *0.020* | *0.038* | *47.20%* | | | | | |
| ***Right*** | *0.030±0.004* | *0.021* | *0.042* | *50.59%* | | | | | |
| **CX** | 0.004±0.001 | 0.001 | 0.007 | 85.13% | F(3,73)=0.727, p=0.539 | 75.33% | 75.30% | 85.13% | 66.70% |
| **OTH** | 0.294±0.065 | 0.087 | 0.408 | 78.65% | F(3,73)=0.919, p=0.436 | 55.55% | 59.48% | 78.65% | 40.37% |

830

105
