## Supplementary material for "Natural variability in bee brain size and symmetry revealed by micro-CT imaging and deep learning": S5 Table

**S5 Table. Correlation between absolute neuropil volumes (bottom left) and between relative neuropil volumes (top right) for bumblebees (N=77).** Pearson correlation coefficient and p-value are given. Strong correlations (r>0.40) and significant correlations (p<0.05) are displayed in bold. Brain areas are labelled using the same abbreviations as in Fig 2.

| 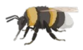 | AL | MB | OL | ME | LO | CX | OTH |
|---|---|---|---|---|---|---|---|
| **AL** | | 0.28 **(p=0.015)** | **0.41 (p<0.001)** | 0.32 **(p=0.004)** | **0.60 (p<0.001)** | -0.03 (p=0.799) | **-0.57 (p<0.001)** |
| **MB** | **0.76 (p<0.001)** | | 0.32 **(p=0.005)** | 0.30 **(p=0.008)** | 0.30 **(p=0.009)** | 0.07 (p=0.541) | **-0.78 (p<0.001)** |
| **OL** | **0.79 (p<0.001)** | **0.85 (p<0.001)** | | **0.99 (p<0.001)** | **0.80 (p<0.001)** | -0.20 (p=0.078) | **-0.82 (p<0.001)** |
| **ME** | **0.74 (p<0.001)** | **0.82 (p<0.001)** | **0.99 (p<0.001)** | | **0.69 (p<0.001)** | -0.18 (p=0.109) | **-0.79 (p<0.001)** |
| **LO** | **0.86 (p<0.001)** | **0.83 (p<0.001)** | **0.91 (p<0.001)** | **0.84 (p<0.001)** | | -0.21 (p=0.061) | **-0.73 (p<0.001)** |
| **CX** | **0.49 (p<0.001)** | **0.52 (p<0.001)** | **0.50 (p<0.001)** | **0.48 (p<0.001)** | **0.49 (p<0.001)** | | 0.06 (p=0.608) |
| **OTH** | **0.57 (p<0.001)** | **0.59 (p<0.001)** | **0.53 (p<0.001)** | **0.48 (p<0.001)** | **0.60 (p<0.001)** | **0.51 (p<0.001)** | |
