## Supplementary material for "Natural variability in bee brain size and symmetry revealed by micro-CT imaging and deep learning": S1 Fig

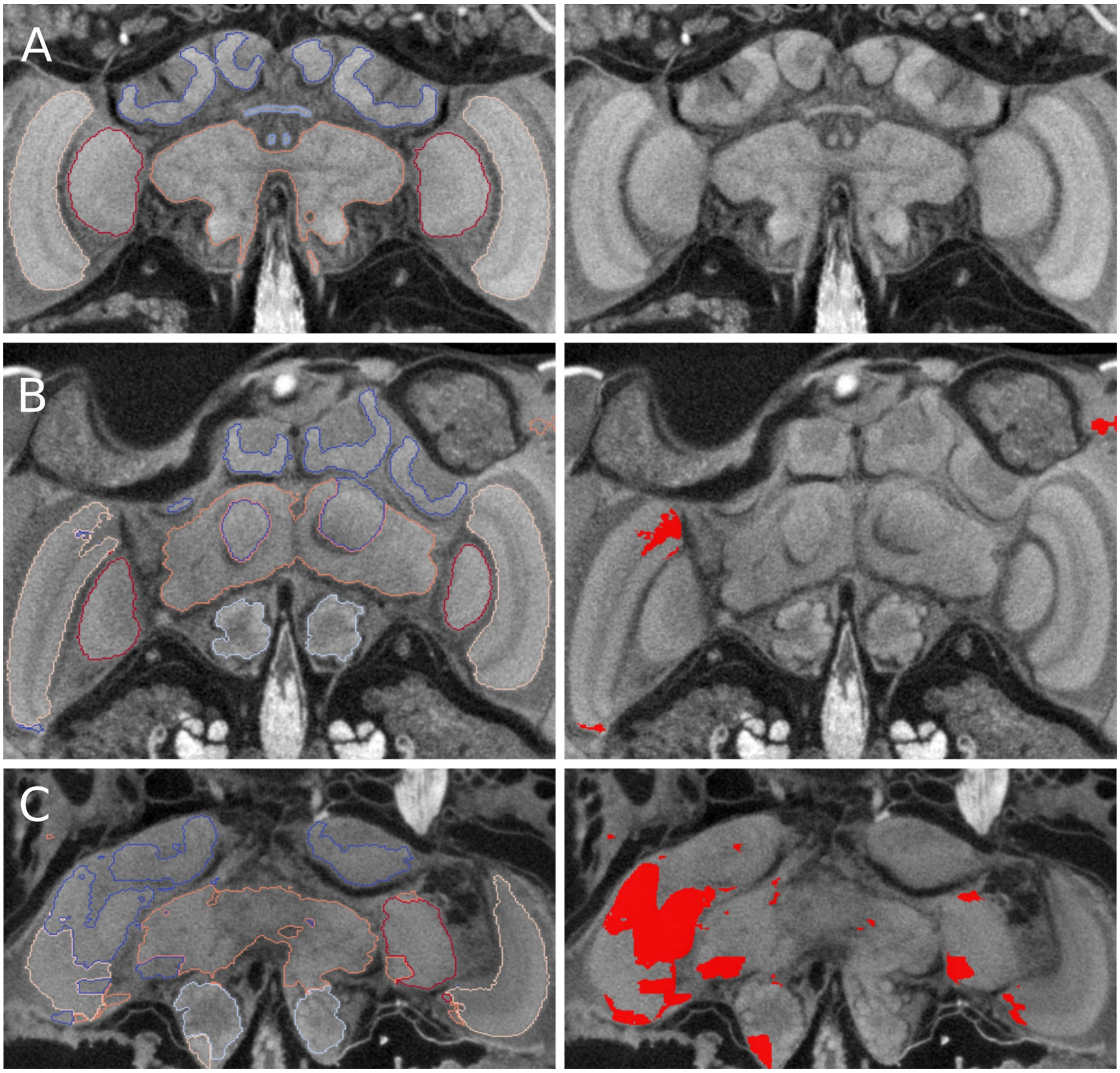


**S1 Fig. Segmentation results (left) and segmentation errors (right) of Biomedisa's deep neural network trained on 26 honey bee CT scans.** (**A**) Correct segmentation without errors (bee ID 87, hive H4). (**B**) Partly flawed segmentation results with a typical outlier on the right edge of the image (bee ID 64, hive H5, segmentation accuracy: ME 97.6%, total 99.1%). (**C**) Significantly flawed segmentation result (bee ID 98, hive H6, total segmentation accuracy 87.7%).
