## Supplementary material for "Natural variability in bee brain size and symmetry revealed by micro-CT imaging and deep learning": S2 Fig

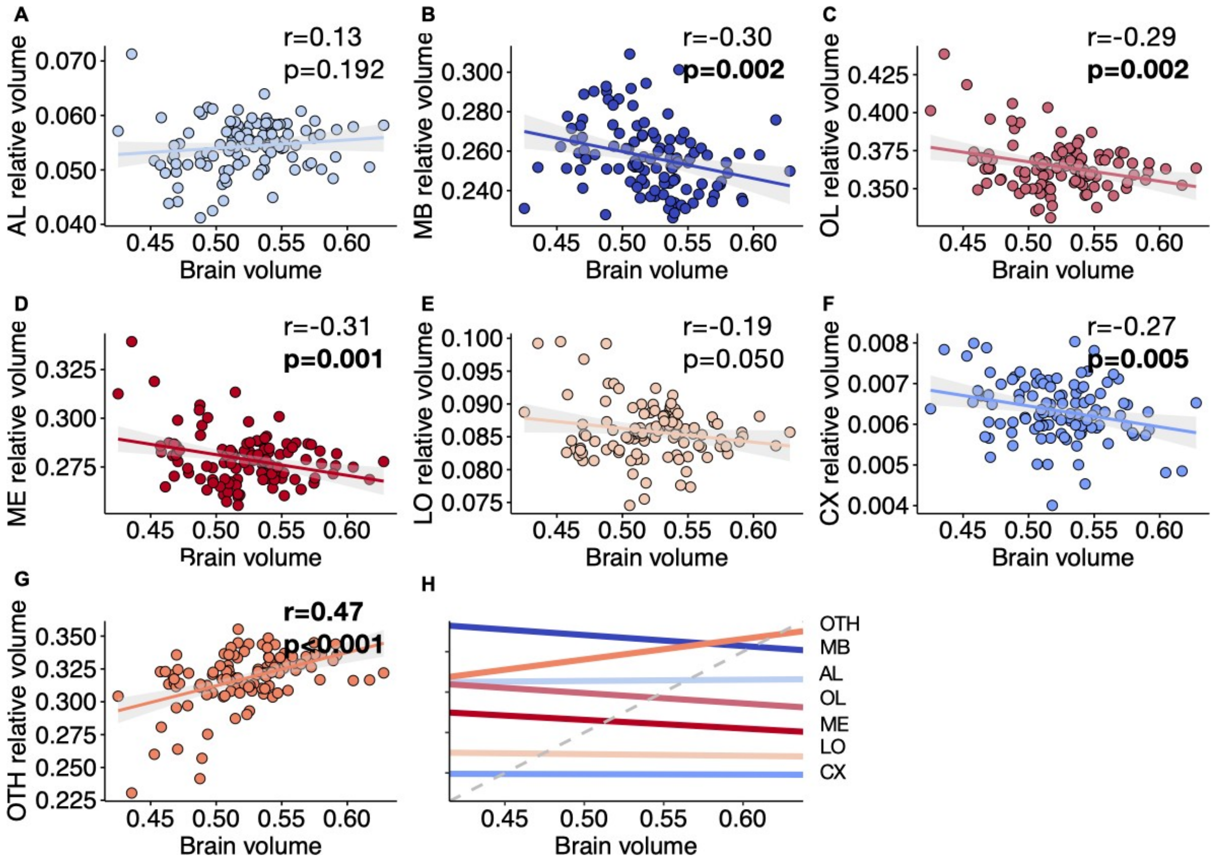


**S2 Fig. Correlation between relative volumes of neuropils and total brain volume (mm$^3$) for honey bees (N=110).** (**A**) Antennal lobes (AL). (**B**) Mushroom bodies (MB). (**C**) Optic lobes (OL). (**D**) Medullae (ME). (**E**) Lobulae (LO). (**F**) Central complex (CX). (**G**) Other neuropils (OTH). Regression lines displayed with 95% confidence intervals. Pearson correlation coefficient (r) and p-value are given. Strong correlations (r>0.40) and significant correlations (p<0.05) are displayed in bold. (**H**) Linear correlations for the different neuropils relative volume (y-axis not given: differs for each neuropil). The grey dashed line indicates true isometric correlation (slope=1).
