## Supplementary material for "Natural variability in bee brain size and symmetry revealed by micro-CT imaging and deep learning": S3 Fig

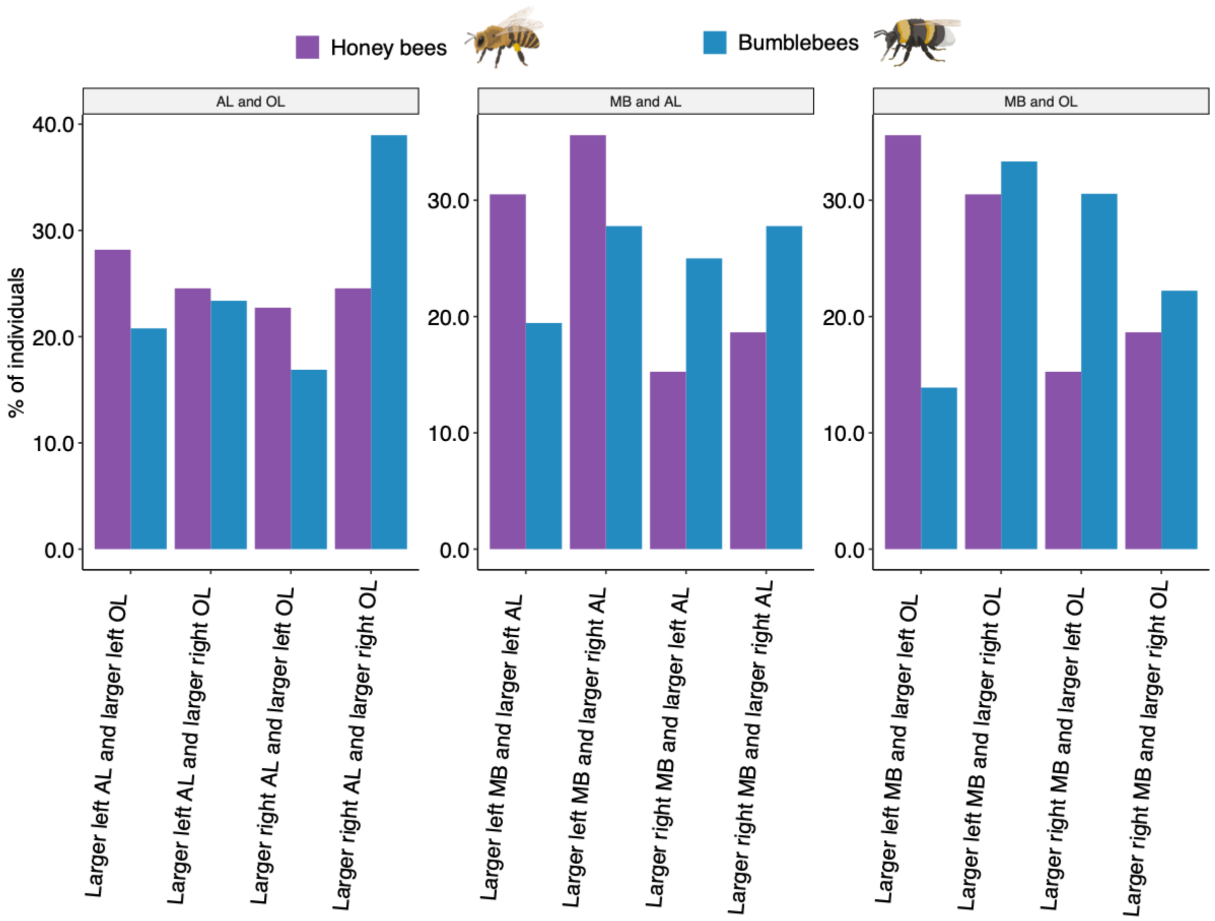


**S3 Fig. Percentage of individuals per asymmetry categories for honey bees (*purple*) and bumblebees (*blue*).** AL and OL (N=110 honey bees, N=77 bumblebees), MB and AL and MB and OL (N=59 honey bees, N=36 bumblebees).
